# Switchable client specificity in a dual functional chaperone coordinates light harvesting complex biogenesis

**DOI:** 10.1101/2024.11.21.624570

**Authors:** Alex R. Siegel, Gerard Kroon, Peter E. Wright, Shu-ou Shan

## Abstract

The proper assembly of light harvesting complexes (LHCs) is critical for photosynthesis and requires the biogenesis of light-harvesting chlorophyll a,b-binding proteins (LHCPs) to be coordinated with the biosynthesis of chlorophylls (Chl). The mechanism underlying this coordination is not well understood. Here we show that a conserved molecular chaperone in chloroplasts, cpSRP43, provides a molecular thermostat that helps maintain this coordination. cpSRP43 undergoes a conformational rearrangement between a well-folded *closed* state and a partially disordered *open* state. *Closed* cpSRP43 is dedicated to the *de novo* biogenesis of LHCPs, whereas *open* cpSRP43 protects multiple Chl biosynthesis enzymes from heat-induced destabilization. Rising temperature shifts cpSRP43 to the *open* state and thus enables it to protect Chl biosynthesis enzymes that are heat-destabilized. Our results reveal the molecular basis of a post-translational mechanism for the thermo-adaptation of LHC biogenesis. They also demonstrate how an ATP-independent chaperone uses conformational dynamics to switch its activity and client selectivity, thereby adapting to different proteostatic demands under shifting environmental conditions.

**Teaser:** A thermo-switchable molecular chaperone helps coordinate light harvesting complex assembly during photosynthesis.

## Introduction

The capture and conversion of sunlight through photosynthesis provides >99% of the energy used by life on earth. To optimize the efficiency of this process, photosynthetic organisms evolved antenna-like complexes that funnel photons into photosynthetic reaction centers. In photosynthetic algae and land plants, the antenna for photosystem II is formed by the light-harvesting complexes, which comprise the LHCP family of proteins bound to chlorophyll (Chl) and other photosynthetic pigments (*1*). LHCPs are integral membrane proteins containing three transmembrane domains and comprise the most abundant family of membrane proteins on earth (*2*). These hydrophobic proteins are nuclear-encoded, synthesized in the cytosol, and highly prone to misfolding and aggregation in the aqueous environments of the cytosol and stroma during their transport to the thylakoid membrane. Thus, the *de novo* biogenesis of LHCPs requires effective chaperone and transport machineries that maintain them in translocation-competent states and mediate their targeted delivery to the thylakoid membrane. The proper folding of LHCPs in the thylakoid membrane further requires the binding of Chl, which are supplied by the tetrapyrrole biosynthesis (TBS) pathway (*3–5*). Reciprocally, the buildup of free Chl or its biosynthetic precursors in the absence of protein binding leads to excess reactive oxygen species (ROS) that can be cytotoxic (*6*). Therefore, proper LHC assembly requires the supply of Chl to be precisely coordinated with the *de novo* biogenesis of LHCPs. The mechanism behind this coordination is not well understood.

cpSRP43, a conserved molecular chaperone that co-evolved with the LHCs, participates in both branches of LHC biogenesis (*7*). cpSRP43 is a subunit of the chloroplast signal recognition particle (cpSRP) responsible for mediating sequence-specific, high affinity recognition of newly imported LHCPs, effectively protecting them from misfolding and aggregation in the chloroplast stroma (*8, 9*). This function is carried out by the substrate binding domain (SBD) comprising four ankyrin repeat motifs (ARMs) capped by an N-terminal chromodomain (CD1) (*8–10*). The other cpSRP subunit, cpSRP54, binds the second chromodomain (CD2) of cpSRP43 via a conserved C-terminal motif (54C) and allosterically enhances cpSRP43’s chaperone activity towards LHCPs (*11–15*). cpSRP further mediates protein-protein contacts at the thylakoid membrane, via the interaction of cpSRP54 with its receptor cpFtsY and the binding of the third chromodomain (CD3) of cpSRP43 to the Alb3 insertase, forming a dedicated pathway for the membrane transport and insertion of the LHCPs (*16–22*). Loss of cpSRP43, cpSRP54, or cpFtsY results in growth retardation, weak pigmentation, and reduced LHCP levels in *Arabidopsis thaliana*, indicating the essential role of the cpSRP pathway in LHC biogenesis (*23–25*).

In addition to its canonical role in LHCP delivery, more recent work found that cpSRP43 also acts as a chaperone to prevent the heat-induced aggregation of multiple enzymes in the TBS biosynthetic pathway (*7, 26*). These biochemically detected chaperone activities were corroborated in studies of *A. thaliana chaos* mutants lacking cpSRP43, which show reduced levels of at least three TBS enzymes including glutamyl tRNA reductase (GluTR), the rate-limiting enzyme in TBS biosynthesis, and GUN4 and CHLH, components of the Mg chelatase complex (*26*). This phenotype is more pronounced during heat stress, when all three enzymes were substantially destabilized, and the levels of Chl and multiple Chl precursors were reduced in *chaos* plants compared to wildtype plants (*26*). The participation of cpSRP43 in both LHCP biogenesis and Chl biosynthesis suggests that it could provide a post-translational mechanism of regulation to balance the levels of LHCPs and Chl during LHC biogenesis (*7*).

However, the molecular basis underlying the potential regulatory role of cpSRP43 remains elusive. It is unclear how a small, ATP-independent chaperone such as cpSRP43 recognizes and protects two distinct types of client proteins, newly synthesized LHCPs and mature TBS enzymes, that differ in size, composition, structure, and folding state. Notably, cpSRP54 enhances the chaperone activity of cpSRP43 towards the LHCPs but represses its activity towards the TBS enzymes (*10, 26*). This observation indicates that only the apo form of cpSRP43 is capable of TBS protection and suggests that cpSRP43 uses distinct molecular mechanisms to recognize the two different classes of clients. The mechanism of this activity switch is unclear, nor how cpSRP43 selects between its two classes of client proteins to balance the supply of LHCP and Chl during LHC assembly.

Early crystal structures showed cpSRP43 in a structured *closed* conformation, in which the helices in all four ARMs are well folded and tightly pack against one another (*9*). However, NMR studies detected a second *open* conformation of cpSRP43 in equilibrium with the *closed* state (*10, 27*). In the *open* state, the two C-terminal ARMs in the cpSRP43 SBD unravel and become partially disordered (*10, 27*). While we previously proposed and provided evidence that the *open* state of cpSRP43 allows this chaperone to turn ‘off’ its activity at the thylakoid membrane during LHCP transport (*10, 27*), the question remains about why a seemingly ‘inactive’ *open* conformation evolved in cpSRP43.

In this work, we define the role of the different conformational states of cpSRP43 using rational mutations and experimental conditions that shift the conformational equilibrium in this chaperone. Biochemical and biophysical studies showed that *closed* cpSRP43 specifically binds and protects LHCPs from aggregation, whereas *open* cpSRP43 is solely responsible for the thermoprotection of TBS enzymes. The *closed*-to-*open* transition is strongly temperature-dependent and, together with the release of cpSRP54, unleashes *open* cpSRP43 to stabilize TBS enzymes at elevated temperatures. We propose a model in which the conformational switch of cpSRP43 provides a molecular thermostat that enables photosynthetic organisms to rapidly respond to rising temperature and maintain coordinated LHC biogenesis.

## Results

### cpSRP43 undergoes a heat-induced conformational change

We previously used ^19^F-NMR to monitor the conformational dynamics of cpSRP43 (*27*). The spectra of cpSRP43 site-specifically labeled with an ^19^F probe, 3-bromo-1,1,1-trifluoroacetone (BTFA), showed two NMR peaks indicating two conformations in slow exchange, which were assigned to a partially disordered *open* state and a rigidly folded *closed* state (*27*). Notably, this conformational change is strongly temperature-dependent: the *closed* state dominated at temperatures below 30 °C, the *open* state became populated between 30 – 40 °C and dominated above 40 °C (Figs. 1A and 1B, red). The heat-induced opening of cpSRP43 could also be detected by circular dichroism (CD), which measures the secondary structure content of proteins (*28*). The CD spectra of wildtype (WT) cpSRP43 changed significantly between 25 °C and 55 °C, with a loss in molar ellipticity (θ) at 222 nm at 55 °C (Fig. 1C), consistent with the partial unfolding of α-helices in the C-terminal ARMs of cpSRP43 in the *open* state (*27*). This conformational change is reversible, with >80% of helical content restored at 25 °C after three heating cycles (Fig. 1C). The temperature dependence of θ_222_ agreed well with that from ^19^F-NMR measurements and yielded a mid-transition temperature (T_m_) of 35 °C (Figs. 1B and 1D, red; Table 1). These results show that elevated temperature induces the *closed*-to-*open* transition of cpSRP43.

**Table 1.** Summary of the mid-transition temperatures of the *open*-to-*closed* conformational change (Tm) for indicated cpSRP43 variants under different conditions.**\***

| [Urea]<br>(M) | FL or<br>SBD | [NaCl]<br>(mM) | R189L | WT | DE3NQ | DE6NQ |
| --- | --- | --- | --- | --- | --- | --- |
| 0 | FL | 0 | < 25† | 29.1 ± 0.11 | 39.9 ± 0.10 | 43.6 ± 0.51 |
|  |  | 50 | < 25† | 35.4 ± 0.13 | 43.3 ± 0.14 | 43.4 ± 0.31 |
|  |  | 100 | < 25† | 37.8 ± 0.09 | 43.5 ± 0.18 | 44.3 ± 0.27 |
|  |  | 200 | 26.4 ± 0.24 | 40.3 ± 0.08 | 45.0 ± 0.17 | 44.7 ± 0.31 |
|  | SBD | 200 | 30.9 ± 0.15 | 41.8 ± 0.14 | 41.8 ± 0.14 |  |
| 0.5 | FL | 200 | < 25† | 38.3 ± 0.11 | 43.2 ± 0.17 |  |
| 1 |  | 200 | < 25† | 35.0 ± 0.16 | 41.1 ± 0.20 |  |

\* Values are represented as fitted values ± standard error of the fits.
† Limits shown when the $T_m$ is below the lowest temperature tested (25 °C) and not measurable.

**Fig. 1.**
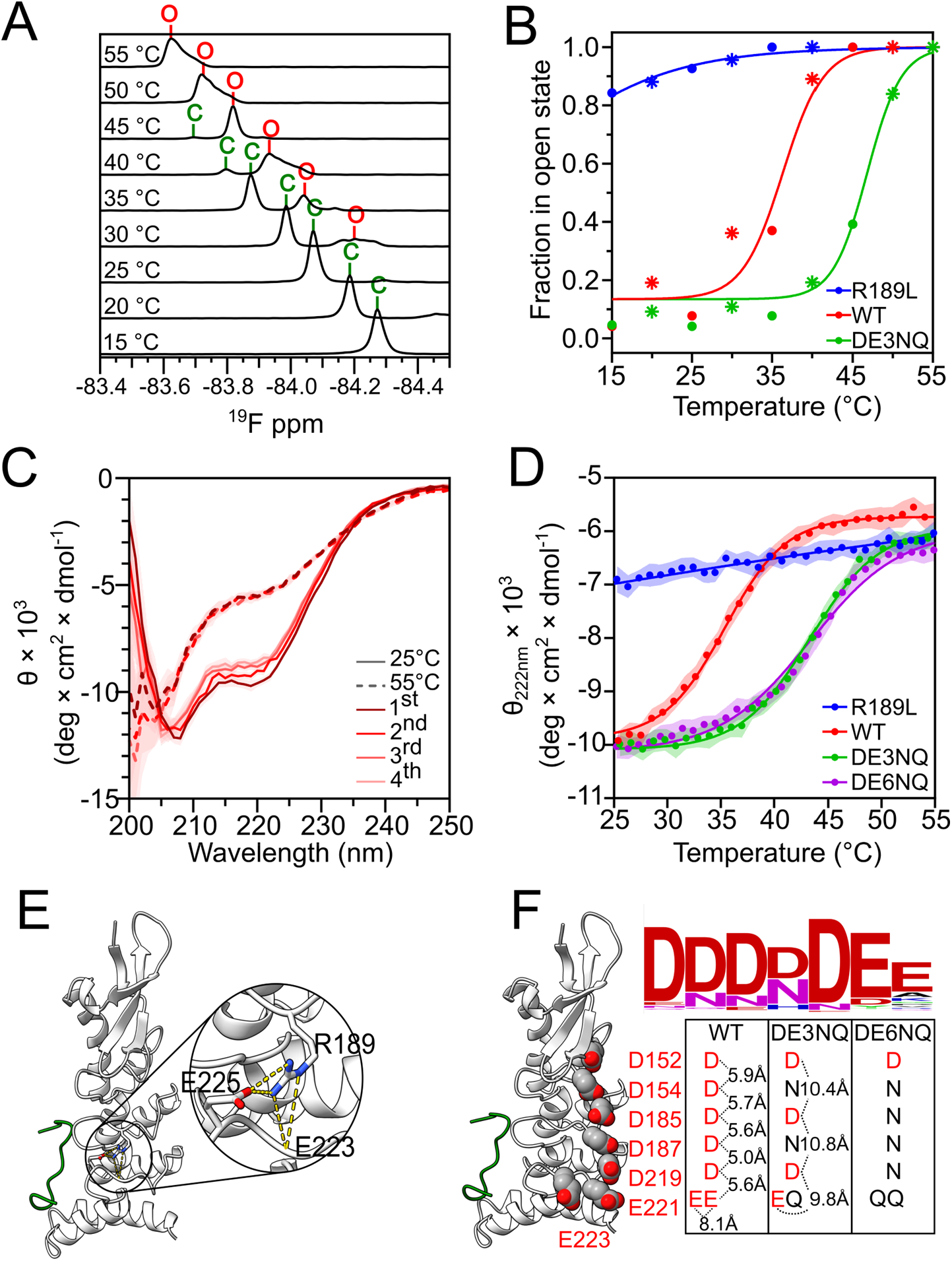
Identification of mutations that stabilize the open or closed state of cpSRP43. (**A**) ^19^F-NMR spectra of BTFA-labeled WT cpSRP43 at the indicated temperatures. The *open* (red ‘O’) and *closed* (green ‘C’) state peaks were assigned in Siegel et al, 2020^***27***^. Spectra were collected in CD buffer with 50 mM NaCl. (**B**) The fraction of cpSRP43 variants in the open state at different temperatures determined by ^19^F-NMR. Data were from the spectra shown in part A and Fig. S1. ‘*’ denotes the data collected after the completion of one heating cycle, where 10-20% of cpSRP43 did not return to the *closed* state. (**C**) CD spectra of WT cpSRP43 at 25 °C (solid) and 55 °C (dashed). Data were collected in CD buffer with 50 mM NaCl and repeated for four heat cycles (dark to light red). Values represent mean residue ellipticity ± S.D. over 5 seconds. (**D**) Temperature dependence of helical content for R189L, WT, DE3NQ, and DE6NQ cpSRP43, measured as per residue molar ellipticity (θ) at 222 nm by CD in CD buffer with 50 mM NaCl. Values represent mean residue ellipticity ± S.D over 10 seconds. The lines are fits of the data to Eq. 2, and the obtained T_m_ values are summarized in Table 1. (**E, F**) *Closed* state structure of the SBD of *A. thaliana* cpSRP43 (PDB: 3DEP^***9***^) showing the hydrogen bonding interactions of R189 (E) or the 7 closely spaced acidic residues (F, left). Sequence conservation of these acidic residues across 1000 land plants are shown in WebLogo (F, upper right).^***47***^ The charge neutralizing mutations are indicated (F, lower right).

To rigorously define the function of cpSRP43 in its different conformations, we sought conservative mutations that would bias the chaperone towards either conformational state. To favor the *open* state, we introduced an R189L mutation that disrupts two hydrogen bonds in the *closed* state structure of cpSRP43 (Fig. 1E) (*9*). ^19^F-NMR and CD measurements showed that, the *open* state is much more populated in R189L compared to WT cpSRP43 (Figs. S1-S2). The estimated T_m_ for opening is 10 – 15 °C for R189L, over 20 °C lower than that for WT cpSRP43 (Fig. 1B, 1D, blue). Thus, the majority of R189L is in the *open* state at room temperature (25 °C) under physiological salt (50 mM NaCl) conditions. The ^19^F-NMR chemical shifts of *open* and *closed* R189L are the same as those of WT cpSRP43 (Fig. S1). In addition, under conditions that stabilize the *closed* state, including low temperature (17 °C), higher salt (see Fig. S6 later), and the presence of the FDPLGL-containing L18 peptide derived from the cpSRP43 recognition motif in LHCP, the ^1^H,^15^N-HSQC spectra of WT cpSRP43 and R189L overlayed well (Fig. S3). Only residues near the FDPLGL binding site or the site of mutation showed significant chemical shift perturbations. These observations strongly suggest that the R189L mutation shifts the conformational equilibrium of cpSRP43 towards the *open* state, without impacting the *closed* state structure.

To drive cpSRP43 towards the *closed* state, we noted that the SBD has an acidic surface with seven evolutionarily conserved Asp and Glu residues in close proximity (Fig. 1F), generating electrostatic repulsion that would destabilize compact folding in the SBD. Charge-neutralizing mutations at these residues in variants DE3NQ (D154/187N, E221Q) and DE6NQ (D154/185/187/219N, E221/223Q) are predicted to stabilize the *closed* state by 3.3 and 3.9 kcal, respectively, at physiological salt concentration (Eq. 3 in Methods (*29*) and Fig. S4, A-B). Consistent with these predictions, the T_m_ value for the *closed*-to-*open* transition was shifted to 43 °C for DE3NQ and DE6NQ, ∼8 °C higher than that for WT cpSRP43 (Figs. 1B, 1D, S1-S2, green and purple; Table 1). The CD spectra and ^19^F NMR chemical shifts of these mutants were the same as WT cpSRP43 at temperatures far below and above their respective T_m_’s (Figs. S1-S2). Except for residues near the mutated sites, most peaks in the ^15^N-HSQC spectra of DE3NQ match the assignments for *closed* cpSRP43 (Fig. S3) (*10*), providing additional evidence that the mutations do not affect the *closed* state structure. Thus, the charge-neutralizing mutations shift the conformational equilibrium of cpSRP43 towards the *closed* state.

To independently verify the effects of the mutations on cpSRP43 conformation, we took advantage of the fact that the partially disordered *open* state is favored by low doses of chemical denaturants (*27*). CD and ^19^F-NMR measurements showed that, compared to WT and R189L cpSRP43, the opening of DE3NQ is more resistant to urea (Fig. S5; Table 1). This provides further evidence for the strong stabilization of the *closed* state by charge-neutralizing mutations on the acidic surface of cpSRP43.

Finally, we explored the effect of salt to further tune the conformation of cpSRP43. Charge screening at higher ionic strength reduces the electrostatic repulsion at the acidic surface of cpSRP43 and is predicted to stabilize the *closed* state by ∼2.3 kcal/mol at 200 mM NaCl relative to 0 mM NaCl (Fig. S4C). CD measurements showed that this was indeed the case: the *closed* state was significantly stabilized at higher salt concentrations in WT cpSRP43 and R189L (Fig. S6; Table 1). In contrast, the salt dependence was much smaller with DE3NQ and negligible in DE6NQ (Fig. S6), as predicted (Fig. S4C). Thus, the *open-*to*-closed* rearrangement can also be tuned by salt concentration in addition to temperature.

Together, the results in this section show that the *open*-to-*closed* transition of cpSRP43 can be regulated by diverse environmental factors including temperature, ionic strength, and low doses of denaturants. Importantly, we successfully isolated rational mutations that modulate this rearrangement, with R189L favoring the *open* state and DE3NQ and DE6NQ hyper-stabilizing the *closed* state, without altering the structure of either conformation.

### Closed cpSRP43 chaperones LHCP

Using the mutations and experimental conditions that drive cpSRP43 towards different conformations, we aimed to define the activity of each conformational state. We first tested the chaperone activity towards its canonical substrate, newly synthesized LHCPs, using an established assay that detects large aggregates based on turbidity at 360 nm (*8*). Dilution of urea-denatured LHCP into buffer led to rapid and extensive aggregation, whereas dilution into buffer containing WT cpSRP43 resulted in quantitative, dose-dependent reductions in LHCP aggregation (Fig. 2A). The chaperone concentration dependence of LHCP solubilization yields solubilization constants (*K*_*sol*_), which represent apparent dissociation constants of the chaperone for LHCP as it competes with the aggregation of the latter. R189L was less effective at chaperoning LHCP compared to WT and DE3NQ at 25 °C, with a >20-fold increase in *K*_*sol*_ (Fig. 2, A-B; Table 2A). This defect was more pronounced at 35 °C and 45 °C, at which little to no solubilization of LHCP was observed even at 16 µM R189L. In contrast, WT cpSRP43 and DE3NQ remained effective at solubilizing LHCP at 35 °C. The high chaperone activity of DE3NQ persisted even at 45 °C, at which WT cpSRP43 displayed weaker chaperone activity, consistent with the higher stability of the *closed* state in DE3NQ than WT cpSRP43 (Fig. 2, A-B; Table 2A). The activity of the cpSRP43 variants towards LHCP across different temperatures correlated with the fraction of chaperone in the *closed* state from CD measurements (Fig. 2C). These data provide definitive evidence that *closed* cpSRP43 is responsible for chaperoning LHCP.

**Table 2.** Summary of the interaction of cpSRP43 variants with LHCP, the L11 peptide, and 54C peptide.*

**A.** Summary of the $K_{sol}$ values ( μM) for LHCP.
| **T (°C)** | **R189L** | **WT** | **DE3NQ** |
| --- | --- | --- | --- |
| 45 °C | N.D.† | $3.5 \pm 1.0$ | $0.025 \pm 0.03$ |
| 35 °C | $62 \pm 18$ | $0.029 \pm 0.01$ | $0.0096 \pm 0.008$ |
| 25 °C | $0.51 \pm 0.09$ | $0.012 \pm 0.01$ | $0.0029 \pm 0.01$ |

**B.** Summary of the $K_d$ values ( μM) for L11 binding to cpSRP43 variants.
| **T (°C)** | **R189L** | **WT** | **DE3NQ** |
| --- | --- | --- | --- |
| 45 °C | $140 \pm 6$ | $0.74 \pm 0.024$ | $0.064 \pm 0.006$ |
| 35 °C | $2.4 \pm 0.09$ | $0.023 \pm 0.002$ | $0.015 \pm 0.002$ |
| 25 °C | $0.19 \pm 0.02$ | $0.018 \pm 0.004$ | $0.015 \pm 0.004$ |

**C.** Summary of the $K_d$ values ( μM) for 54C binding to cpSRP43 variants.
| **T (°C)** | **R189L** | **WT** | **DE3NQ** |
| --- | --- | --- | --- |
| 45 °C | $7.7 \pm 0.1$ | $5.1 \pm 0.06$ | $3.5 \pm 0.06$ |
| 35 °C | $1.0 \pm 0.02$ | $0.19 \pm 0.005$ | $0.30 \pm 0.004$ |
| 25 °C | $0.083 \pm 0.003$ | $0.029 \pm 0.002$ | $0.051 \pm 0.001$ |

\*All values were measured at 50 mM NaCl and are represented as fitted values ± standard error of the fits.
† Not detectable.

**Fig. 2.**
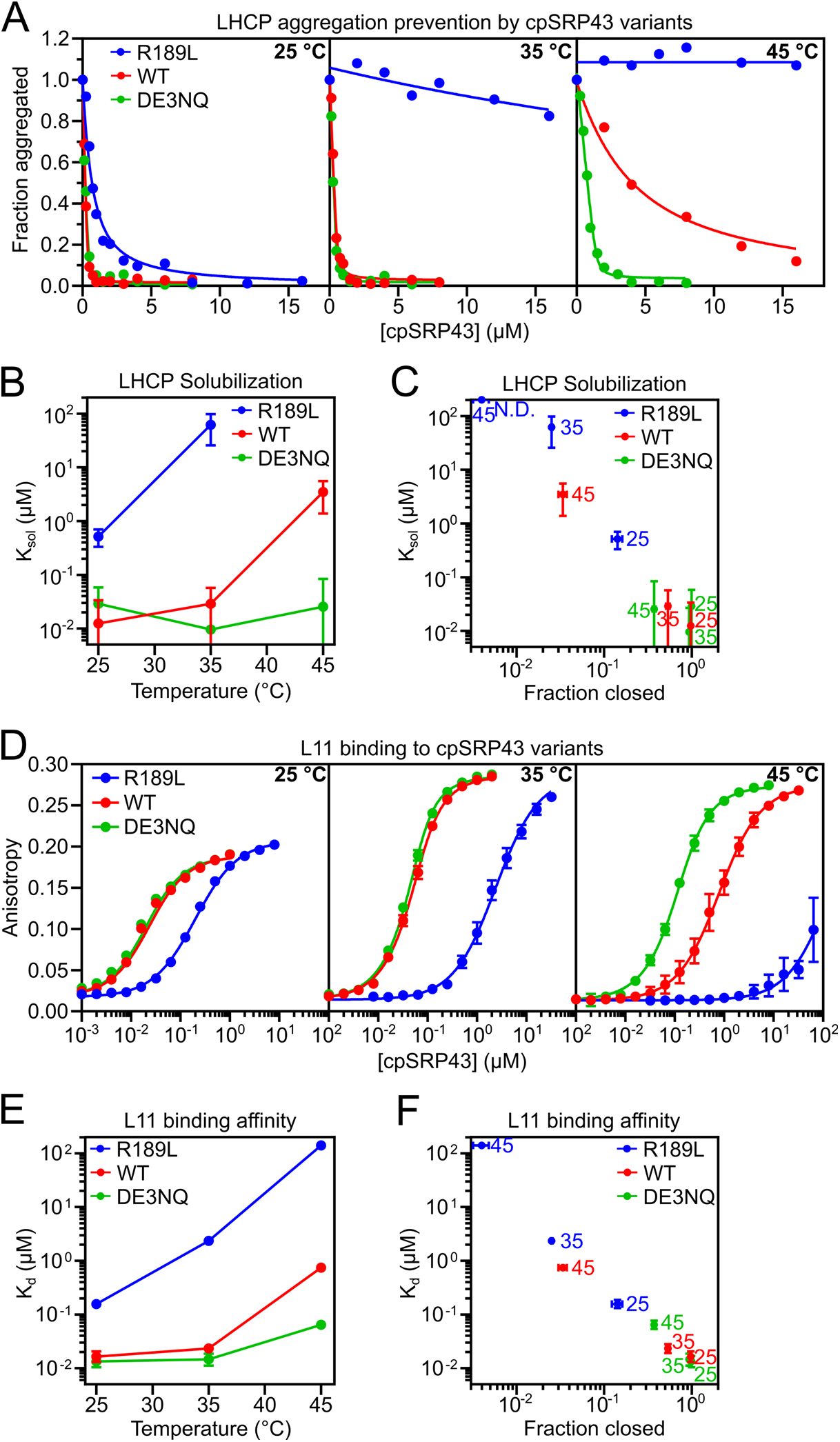
Closed cpSRP43 mediates chaperone activity towards the LHCPs. (**A**) Turbidity assays to measure the protection of LHCP by increasing concentrations of cpSRP43 variants at 25 °C (left), 35 °C (middle), and 45 °C (right) in CD buffer with 50 mM NaCl. Optical density at 360 nm (A360) were normalized to that without cpSRP43. Lines are fits of the data to Eq. 4. (**B, C**) Obtained *K*_*sol*_ values for LHCP were plotted as a function of temperature (B) or fraction of chaperone in the *closed* state (C). Values represent fitted value ± 95% CI of the fit. (**D**) Equilibrium titrations to measure the binding of the indicated cpSRP43 variants to HiLyte-Fluor488-labeled L11 peptide at 25 °C (left), 35 °C (middle), and 45 °C (right) in CD buffer with 50 mM NaCl. Values represent mean ± S.E. for n = 5 reads. Lines are fits of the data to Eq. 6. (**E, F**) Obtained *K*_*d*_ values for L11 peptide binding were plotted as a function of temperature (E) or fraction of chaperone in the *closed* state (F). Values represent fitted value ± 95% CI of the fit. The numbers next to the data points in (C) and (F) indicate the temperatures at which the measurement was made. The fraction of chaperone in the *closed* state was from the measurements in Figure 1D.

Recognition and protection of LHCP by cpSRP43 requires a sequence-specific interaction with the conserved FDPLGL motif in a stromal loop of LHCPs (*9–11, 30, 31*). To test whether high affinity recognition of this motif also occurs exclusively in *closed* cpSRP43, we measured the binding affinity of the cpSRP43 variants for an L11 peptide containing the FDPLGL motif based on cpSRP43-induced anisotropy change of fluorescein-labeled L11 (Fig. 2D). The effects of cpSRP43 mutations and temperature on L11 binding affinity mirrored those on LHCP solubilization by this chaperone (Fig. 2, D-E; Table 2B). The equilibrium dissociation constant (*K*_d_) for L11 binding across the chaperone variants and temperatures also strongly correlated with the fraction of chaperone in the *closed* state (Fig. 2F). Thus, the *closed* state of cpSRP43 specifically recognizes the FDPLGL motif in LHCP and protects this family of membrane proteins from aggregation, whereas the *open* state does not.

### Open cpSRP43 mediates thermoprotection of TBS enzymes

We next asked how the chaperone activity of cpSRP43 towards its second class of clients, the TBS enzymes, depends on its conformational state. Mature, folded GUN4, a model TBS enzyme substrate for cpSRP43 (*26*), aggregates rapidly upon heat treatment as monitored by increased turbidity at 360 nm (Fig. S7). The aggregation time courses of GUN4 fit well to first-order kinetics, and the observed rate constant of aggregation was independent of GUN4 concentration (Fig. S7, A-B; Table S1A). As protein misfolding/unfolding is concentration-independent whereas aggregation is strongly concentration-dependent, these results indicate that GUN4 undergoes rate-limiting misfolding followed by rapid aggregation under heat stress. The misfolding and aggregation of GUN4 is accelerated by rising temperature, plateauing at a rate constant of 0.13 min^-1^, or half-time of 5.3 min, above 42 °C (Fig. S7, C-D; Table S1B).

To assess the ability of cpSRP43 variants to prevent heat-induced GUN4 aggregation, we chose a condition at the midpoint of GUN4 aggregation kinetics (5 min at 42 °C). Under these conditions, the *open* state population varied significantly across the cpSRP43 mutants: 96% for R189L, 65% for WT, and 25% for DE3NQ (Fig. 1D). The cpSRP43-mediated thermoprotection of GUN4 was strongest for mutant R189L, which is primarily in the *open* state under these conditions, and lowest for mutant DE3NQ, which is hyper-stabilized in the *closed* state (Fig. 3A; Table 3A). The same trend was observed for the thermoprotection of a second TBS enzyme, GluTR, by the cpSRP43 variants (Fig. 3B; Table 3B).

**Table 3.** Summary of the chaperone activities of cpSRP43 variants towards TBS enzymes.

**A.** Summary of the $K_{sol}$ values ( μM) for GUN4.\*
| **[NaCl (mM)]** | **FL or SBD** | **R189L** | **WT** | **DE3NQ** |
| --- | --- | --- | --- | --- |
| 50 | FL | $0.079 \pm 0.06$ | $0.25 \pm 0.08$ | $3.2 \pm 0.2$ |
| 50 | SBD | $2.5 \pm 0.4$ | $6.7 \pm 0.4$ | N.D.† |
| 100 | FL | $0.069 \pm 0.09$ | $1.6 \pm 0.3$ | $51 \pm 11$ |
| 200 | FL | $0.035 \pm 0.2$ | $9.8 \ddagger$ | N.D.† |

\* Values were measured at 50 mM NaCl, 42 °C and are shown as fitted value ± standard error of the fits.
† Not Detectable.

‡ $K_{1/2}$ value. **B.** Summary of the $K_{sol}$ values ( μM) for GluTR.\*
| **FL or SBD** | **R189L** | **WT** | **DE3NQ** |
| --- | --- | --- | --- |
| FL | $0.010 \pm 0.03$ | $0.22 \pm 0.1$ | $1.7 \pm 0.6$ |
| SBD | $0.24 \pm 0.09$ | $10 \pm 2$ | N.D.† |

\*Values were measured at 200 mM NaCl, 42 °C and are shown as fitted value ± standard error of the fits.
† Not Detectable.

**Fig. 3.**
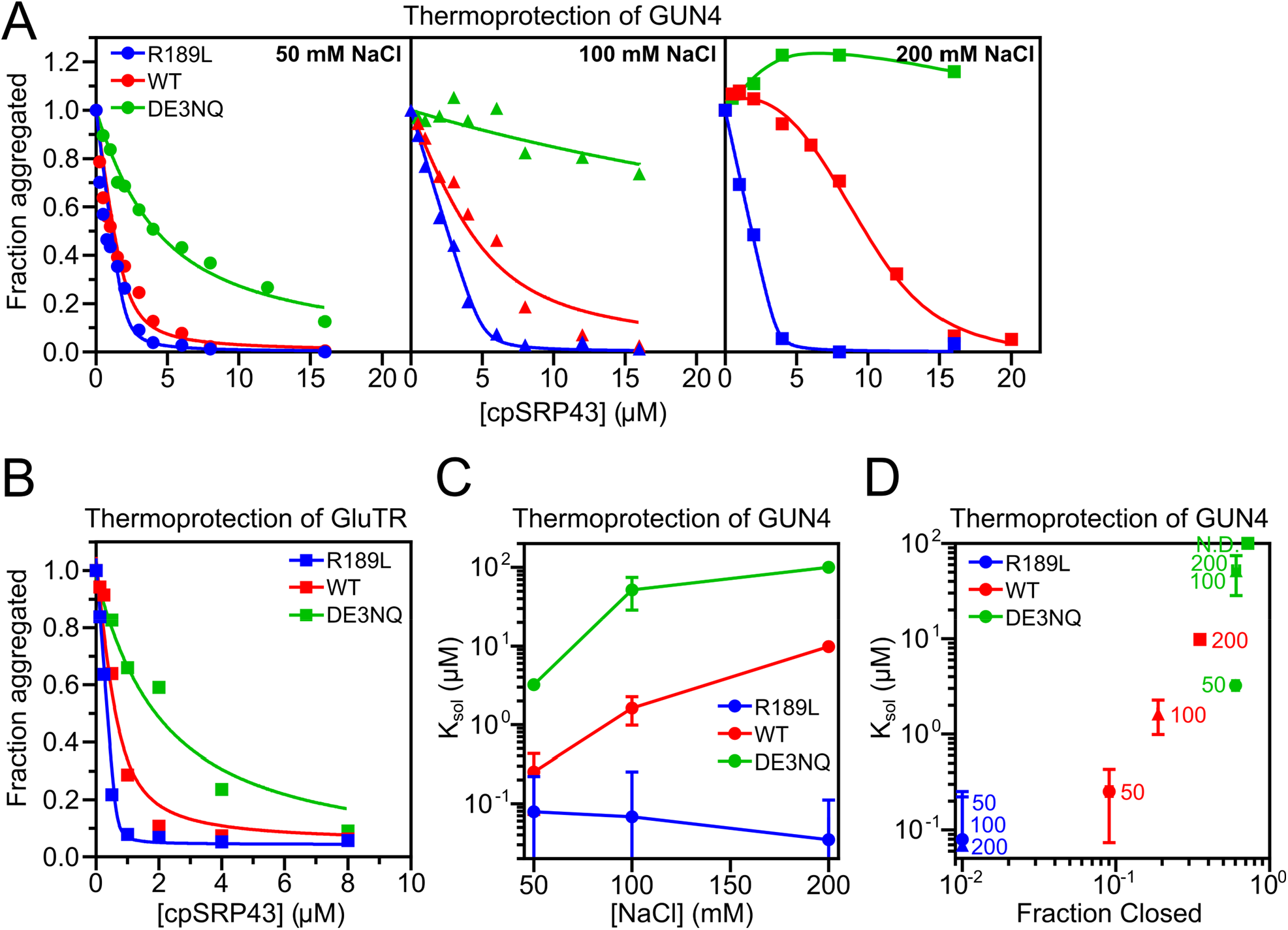
Open cpSRP43 mediates the thermoprotection of TBS enzymes. (**A**) Turbidity assays to measure the heat-induced aggregation of 10 μ*M* GUN4 and its protection by the indicated cpSRP43 variants in LS buffer with 50 (left), 100 (middle), and 200 (right) mM NaCl. Optical density at 360 nm were normalized to that without cpSRP43. The lines are fits of the data to Eq. 4 or to a smoothing spline. (**B**) Same as (A), except with 2.7 µM GluTR as the client protein. (**C, D**) The fitted *K*_*sol*_ values or estimated *K*_*½*_ values from the data in A (see Methods) are plotted as a function of NaCl concentration (C) and fraction of chaperone in the *closed* state (D). The numbers next to the data indicate the NaCl concentration at which the measurements were made. Values represent fitted value ± 95% CI of the fit.

To further tune the conformational equilibrium of cpSRP43 at constant temperature, we increased ionic strength, which reduces the *open* state population (Fig. S6). WT cpSRP43 and DE3NQ showed reduced thermoprotection of GUN4 at higher ionic strength, with a >10-fold rise in *K*_*sol*_ values at 200 mM NaCl compared to 50 mM NaCl (Figs. 3A, 3C; Table 3A). This is consistent with the diminished *open* state population at higher salt in these chaperone variants (Fig. S6). In contrast, R189L, which predominantly populates the *open* state at 42 °C even at 200 mM NaCl, retained effective GUN4 protection (Figs. 3A, 3C, blue; Table 3A). We note that heat-induced aggregation of GUN4 is slower at higher ionic strength (Fig. S7, E-F); hence, the reduced protection of GUN4 by WT cpSRP43 and DE3NQ under these conditions cannot be attributed to more aggressive GUN4 aggregation. The *K*_sol_ values for cpSRP43 variants at different salt concentrations correlated with the population of the chaperone in the *open* state observed by CD (Fig. 3D). Thus, while the *open* state of cpSRP43 lacks activity towards the LHCPs, it is responsible for protecting mature TBS enzymes from heat-induced misfolding and aggregation. In contrast, the *closed* state does not recognize and protect this second class of client proteins.

### cpSRP54 and C-terminal chromodomains directly regulate TBS thermoprotection

We previously found that binding of the other subunit of cpSRP, cpSRP54, weakens with rising temperature (Ji et al 2021 (*26*); Fig. 4, A and B), which correlated with its release from cpSRP43 upon heat stress in *A. thaliana*. However, cpSRP54 also stabilizes the *closed* state of cpSRP43 (*10*) (Fig. S8), the population of which diminishes at elevated temperature. This raises the question: is the heat-induced release of cpSRP54 from cpSRP43 due to the intrinsic temperature sensitivity of their interaction, or to the opening of cpSRP43 at high temperature? To distinguish between these models, we leveraged the set of cpSRP43 variants that opens at different temperatures. cpSRP54 binding was measured based on changes in the fluorescence anisotropy of a fluorescein-labeled 54C peptide bearing the cpSRP43-binding motif of cpSRP54 (*13*). WT cpSRP43 and DE3NQ bound 54C with 3-4 fold higher affinities than R189L, consistent with the higher population of *open* state in R189L and the preference of cpSRP54 for the *closed* state.

**Fig. 4.**
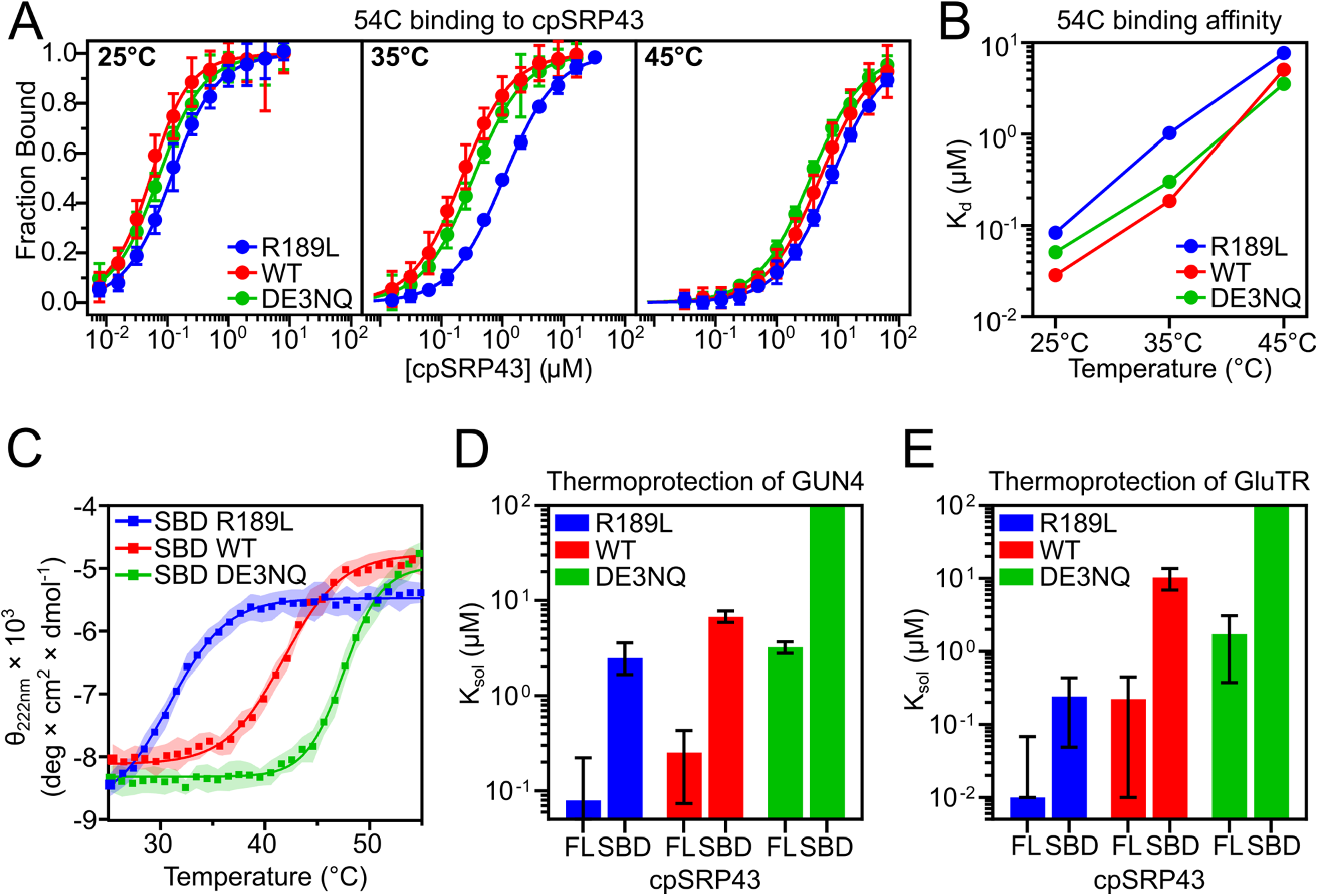
cpSRP54 and C-terminal chromodomains modulate TBS thermoprotection by open cpSRP43. (**A, B**) Equilibrium titrations to measure the binding affinity of the 54C peptide for cpSRP43 variants at rising temperatures (A). Reactions were carried out in CD buffer with 50 mM NaCl. Values represent mean ± S.E. for n = 5 reads. The lines are fits of the data to Eq. 6. The obtained *K*_*d*_ values are plotted as a function of temperature (B). (**C**) Temperature dependence of the helical content for R189L, WT, and DE3NQ cpSRP43(SBD), measured as per residue molar ellipticity (θ) at 222 nm by CD in CD buffer with 200 mM NaCl and fit as in Figure 1D. Values represent mean residue ellipticity ± S.D. of the signal over 10 seconds. (**D, E**) Comparison of the *K*_sol_ values between FL and SBD constructs in the thermoprotection of GUN4 (D) and GluTR (E) for cpSRP43 variants. *K*_*sol*_ values were from fits of the data in Fig. S9 to Eq. 4 and are shown as fitted value ± 95% CI of the fit.

However, the interaction of all three variants with 54C displayed a strong temperature dependence that was largely independent of their conformational equilibrium (Fig. 4, A and B; Table 2C). For example, DE3NQ is predominantly *closed* between 25 °C and 35 °C, over which its 54C binding weakened 6-fold. Similarly, R189L is exclusively *open* between 35 and 45 °C, over which its 54C binding weakened 8-fold (Fig. 4, A-B; Table 2C). These results show that the intrinsic temperature sensitivity of the cpSRP43-cpSRP54 interaction is primarily responsible for the disassembly of cpSRP at high temperature, whereas the conformational state of the cpSRP43 SBD contributes modestly.

While the cpSRP43 SBD is necessary and sufficient to chaperone the LHCPs (*10*), at least one of the C-terminal chromodomains is further required for cpSRP43 to effectively protect TBS enzymes (*26*). However, the presence of CD2 also biases cpSRP43 towards the *open* state (*10*). To distinguish whether the requirement for CD2 is due to its effect on SBD conformation or to a direct interaction with TBS proteins, we examined the impact of deleting CD2 and CD3. For R189L, both the full-length chaperone and the SBD are predominantly *open* at 42 °C (Fig. 4C and Fig S9A; Table 1). If CD2CD3 exerts its effect by enhancing the *open* state population, their deletion should have no impact on the activity of R189L. In contrast to this prediction, R189L(SBD) showed 30-fold reduced chaperone activity towards GUN4 (Fig. 4D and Fig S9B, blue; Table 3A) and 50-fold reduced chaperone activity towards GluTR (Fig. 4E and Fig S9C, blue; Table 3B) compared to full-length R189L, indicating a direct involvement of CD2 and/or CD3 in the thermoprotection of TBS enzymes. In addition, the effect of CD2CD3 deletion on the chaperone activity towards TBS enzymes was consistent across the cpSRP43 variants irrespective of their conformational equilibria (Fig. 4D and E, Fig S9B and C; Table 3). Therefore, the C-terminal chromodomains of cpSRP43 directly participate in the recognition and protection of TBS enzymes under heat stress.

## Discussion

The biogenesis of LHCs is a rate-limiting step in biomass production from solar energy by photosynthetic organisms and presents multiple challenges to the cellular proteostasis network. A particular challenge is the requirement for precise coordination between the biogenesis of LHCPs, mediated by the cpSRP pathway, and the supply of Chl molecules, synthesized by the TBS pathway. Inadequate balance between the two branches leads to the accumulation of aggregated LHCPs or toxic ROS, generating proteostatic or phototoxic stress (*6*). This balance must be further maintained under different environmental conditions, such as varying light intensity and temperature during the day-night cycle and across seasons. Our previous work uncovered chaperone activities of cpSRP43 towards client proteins in both the LHCP biogenesis and Chl synthesis pathways, but how it coordinates the two pathways was unclear. This work elucidates the molecular basis of this coordination and posits cpSRP43 as a molecular thermostat, potentially helping photosynthetic organisms balance the supply of Chl with LHCP biogenesis at elevated temperature.

Our previous biophysical studies demonstrated the presence of two conformations of cpSRP43 at equilibrium, a structured *closed* state and a partially disordered *open* state. Here, we found that this conformational change provides the fundamental mechanism that explains both the dual chaperone activity of cpSRP43 and its regulation by environmental factors. Using rational mutations and conditions that tune this conformational equilibrium, we show that hyperstabilizing cpSRP43 in the *closed* state enhanced chaperone activity towards LHCP but abolished its activity towards two TBS enzymes (Figs. 2-3). The opposite was observed with chaperone variants that populate the *open* conformation (Figs. 2-3). These results provide definitive evidence that each conformational state in cpSRP43 is used to handle a distinct class of client proteins. *Closed* cpSRP43 is dedicated to the chaperoning and membrane transport of newly synthesized and imported LHCPs, whereas *open* cpSRP43 protects mature TBS enzymes from heat stress. Moreover, this conformational change is exquisitely sensitive to temperature, with cpSRP43 transitioning almost completely from the *closed* to the *open* state between 30 – 40 °C, with a mid-transition temperature of 35 °C. This temperature dependence correlates with the acceleration in the misfolding and aggregation of GUN4 over the same temperature range, which became kinetically significant above 34 °C (*k* ∼ 0.008 min^-1^). Thus, this temperature dependent conformational switch enables cpSRP43 to sense and respond to heat stress, unleashing its thermoprotection activity for TBS enzymes precisely when it is needed.

In addition to the conformational change in the cpSRP43 SBD, the C-terminal chromodomains of this chaperone and its binding partner cpSRP54 further modulate chaperone activity towards the TBS enzymes. While the C-terminal chromodomains were previously shown to favor the opening of cpSRP43, the results here provide evidence that they directly contribute to the binding and protection of TBS enzymes. On the other hand, cpSRP54 favors the *closed* conformation of cpSRP43, inhibits its TBS protection activity, and is released from this chaperone at elevated temperatures. We find that the strong temperature dependence of the cpSRP54-cpSRP43 interaction occurs largely independently of its effect on the *open*-to-*closed* transition in the SBD and likely provides a second molecular mechanism to sense rising temperature. Together with the *opening* of the cpSRP43 SBD, the release of cpSRP54 enforces the switch in the client preference of cpSRP43 from the LHCPs to the TBS enzymes at elevated temperature.

We propose a model for how the dual chaperone functions of cpSRP43, enabled by its conformational switch, help coordinate Chl synthesis with LHCP biogenesis for plants experiencing elevated temperature (Fig. 5). At low temperature (< 30 °C for *A. thaliana*), cpSRP43 is predominantly in the *closed* state and tightly bound to cpSRP54, providing a dedicated pathway for the protected transport and insertion of newly imported LHCPs at the thylakoid membrane. Under these conditions, TBS enzymes are stably folded and mediate the biosynthesis of Chl that is integral to LHCP folding and assembly. As temperature rises from 30 to 40 °C, TBS enzymes are increasingly misfolded, aggregated, and susceptible to proteolytic degradation. Over this temperature range, an increasing population of cpSRP43 is released from cpSRP54 and adopts the *open* conformation, which stabilizes TBS enzymes. This allows plants to rapidly respond to rising temperature and maintain the balance of Chl synthesis with LHCP biogenesis.

**Fig. 5.**
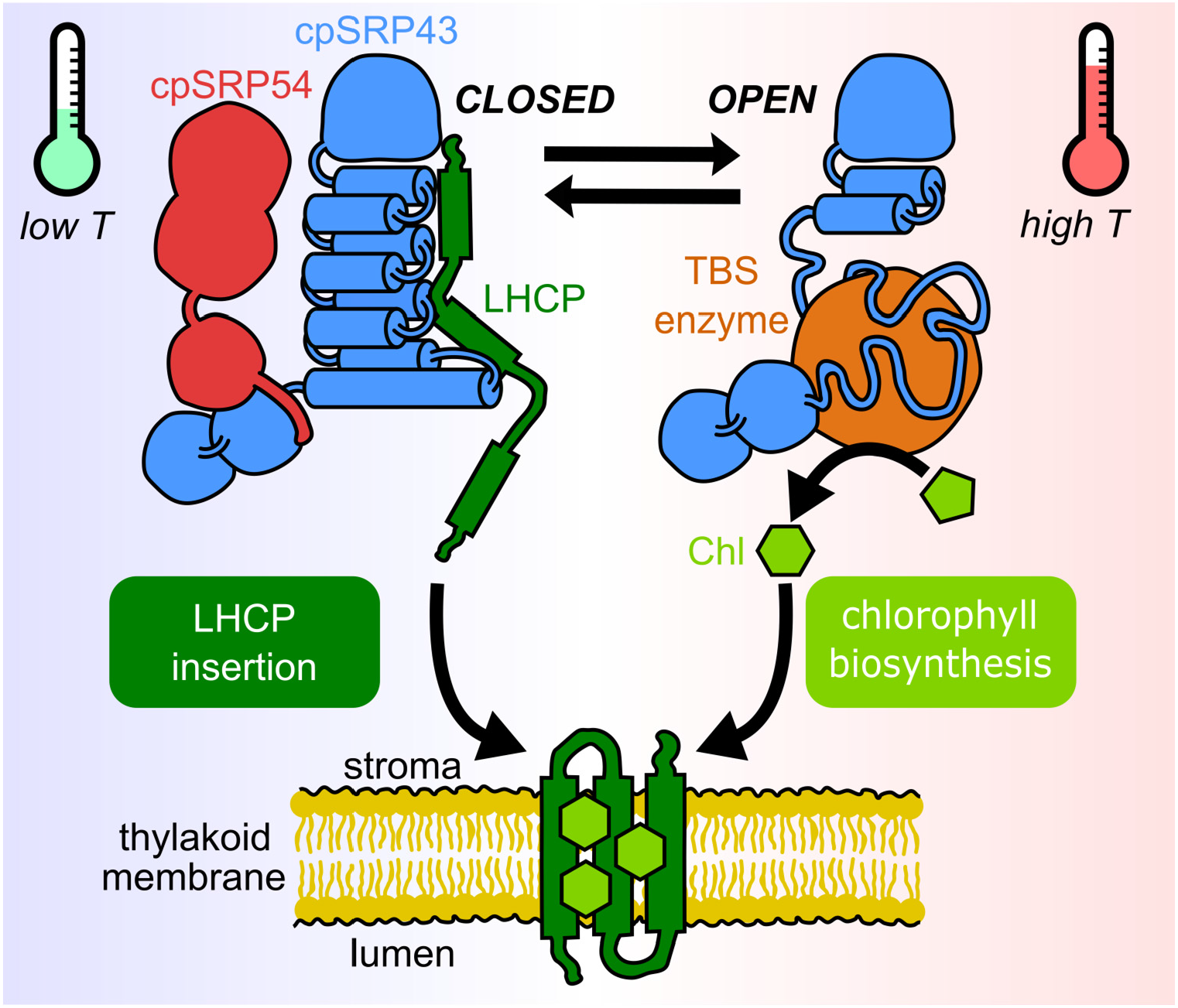
Model for cpSRP43’s dual chaperone activities. Under normal conditions, cpSRP43 primarily populates the *closed* conformation in which its ankyrin repeat motifs in the SBD are tightly folded and further stabilized by interactions with cpSRP54 (left). *Closed* cpSRP43 confers high affinity recognition and protection of newly imported LHCP but has no activity towards TBS enzymes. A fraction of cpSRP43 samples the *open* conformation in which the C-terminal ankyrin repeat motifs become partially disordered (right). *Open* cpSRP43 does not recognize LHCP, but effectively protects mature TBS enzymes from misfolding, aggregation, and resulting proteolytic degradation. Elevated temperature and increased levels of TBS enzymes drive the transition of cpSRP43 to the *open* state, unleashing its thermoprotection activity under conditions where this activity is needed.

We speculate that this mechanism is most relevant during peak sunlight hours in the early afternoon, especially in summertime, during which leaf temperature can rise to ∼35 °C in well hydrated plants. The transition of a fraction of cpSRP43 to the *open* conformation under these conditions provides plants short-term thermoprotection during this modest daily heat stress. In addition, the expression of TBS enzymes is transcriptionally upregulated after sunrise; the increased abundance of these enzymes during daylight could shift a fraction of cpSRP43 to the *open* state, which recognizes the TBS clients, via the law of mass action. These hypotheses remain to be tested *in vivo*. In addition, the physiological role of this mechanism for photosynthetic organisms experiencing long-term or extreme heat stress conditions remains unclear. Finally, our biochemical data suggest that other factors, such as salt concentration and low doses of denaturants, extensively modulate the conformational transition of cpSRP43; whether this conformational change could also respond to additional environmental stresses, such as salinity, remains to be tested.

Our results here provide a novel paradigm in which a molecular chaperone completely alters its activity and client specificity in response to a change in environmental conditions. While temperature (*32, 33*), pH (*34–37*), and oxidative stress (*38, 39*), have all been reported to induce conformational changes in various small, ATP-independent chaperones, in most cases these chaperones switch between an inactive “storage” mode and an active client-binding mode in response to environmental stress. A notable exception is DegP in bacteria (*40*) and its eukaryotic homolog HtrA2 in the mitochondrial intermembrane space (IMS) (*41, 42*), which acts as a chaperone at low temperature and as a protease for the same clients at high temperature. cpSRP43 is unique in that a single chaperone harbors two distinct conformational states that are *each* active towards *different* clients. By regulating the population of its two conformations over a broad range of physiological temperatures, cpSRP43 post-translationally balances the levels of its clients to meet the changing needs for coordinated LHC assembly during light capture. Given that its change in activity arises from an order-to-disorder conformational transition, cpSRP43 also provides an excellent model to understand the mechanisms by which structured versus disordered chaperones specialize in the recognition and protection of distinct types of client proteins.

## Materials and Methods

### Protein expression and purification

Mutations of cpSRP43 were constructed using the QuikChange Mutagenesis procedure (Stratagene) according to manufacturer’s instructions. WT and mutant cpSRP43, LHCP, GUN4, and GluTR were overexpressed and purified as previously described (*7, 26, 43*).

### Circular dichroism

Spectra were collected on an Aviv Model 410 Circular Dichroism Spectrophotometer in 1 mm thick quartz cuvettes containing 300 μL of 10 μM cpSRP43 in CD buffer (20 mM Na_2_HPO_4_, pH 7.4) with 0, 50, 100, or 200 mM NaCl and supplemented with 0.5 or 1 M urea where indicated. Spectra were acquired with 5 seconds averaging in 1 nm increments from 250 to 200 nm at either 25 °C or 55 °C. Thermal melts were acquired by increasing the temperature in 1 °C increments from 25 °C to 55 °C and recording the ellipticity at 222 nm averaged over 10 seconds. Raw ellipticity at 222 nm in instrument units (millidegrees) was converted to per residue molar ellipticity (θ) using Eq 1:

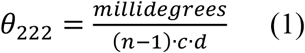

in which *n* is the number of amino acids in the protein (329 for FL, 222 for SBD), *c* is the concentration in *M* (1×10^−5^ *M* or 10 μ*M*), and *d* is the pathlength in mm (1 mm). The temperature dependence of θ_222_ was fit to Eq. 2,

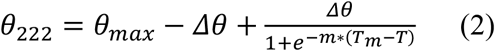

in which *θ*_*max*_ is the molar ellipticity of the open state, *Δθ* is the difference in molar ellipticity between the *open* and *closed* states, *m* is the slope of the sigmoid, *T*_*m*_ is the transition temperature at which the *open* and *closed* states are equally populated, and *T* is the temperature of the measurement.

### Free energy calculation of the effects of mutations and salt

Distances between all pairs of negatively charged residues on the acidic surface (D152, D154, D185, D187, D219, E221, E223) were calculated in ChimeraX based on the structure of *closed* cpSRP43 SBD (PDB: 3DEP(*9*)). The coulombic contribution to the free energy of the *closed* state (*ΔG*_*electric*_) of each pair was computed using Eq. 3a and 3b (*29*) and summed over all pairs.

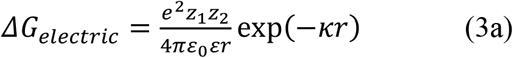

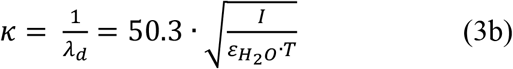

In Eq 3, *e* is the charge of an electron, z_1_ and z_2_ are the charges of the two residue pairs, *ε*_*0*_, *ε*_*H2O*_, and *ε* are the permittivity of free space in a vacuum, in water (estimated to be 78.5), and at the protein surface in water (estimated to be 4), respectively, *κ* is the inverse of Debye length (λ_d_) as defined in Eq 3b, *I* is the ionic strength of buffer and is 0, 0.1, 0.15, and 0.25 for buffers with 0, 50 mM, 100 mM, and 200 mM NaCl, respectively, and *T* is the temperature in K (*29*).

### Chaperone activity

The chaperone activity of cpSRP43 towards LHCP was measured as described previously (*8, 30*). cpSRP43 solution was clarified by ultracentrifugation in a TLA-100 rotor (Beckman Coulter) at 100,000 rpm for 30 min at 4 °C prior to the experiment. LHCP aggregation was initiated by addition of 2 µL of 50 µM LHCP denatured in Urea buffer (8 M urea, 10 mM Tris, 100 mM Na_2_HPO_4_, pH 8.0) to 100 µL of CD buffer with 50 mM NaCl and containing indicated concentrations of cpSRP43 and/or urea. Samples were incubated at the 25, 35, or 45 °C in a water bath for 5 minutes, followed by 5 minutes on ice. The endpoint optical density was recorded at 360 nm on a UV-Vis spectrometer (Beckman Coulter) after equilibrium has been reached (5 minutes). The fraction of aggregated LHCP or TBS enzyme client at equilibrium, *f*, was determined as a function of cpSRP43 concentration ([cpSRP43]) and fit to Eq. 4,

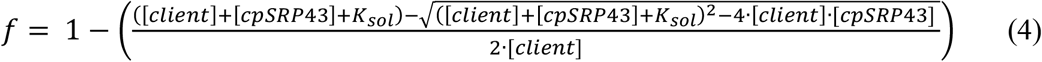

in which [client] is the concentration of client protein, and *K*_*sol*_ is the apparent solubilization constant.

Thermoprotection of TBS enzymes by cpSRP43 was measured as described (*26*). Purified, folded recombinant GUN4 and GluTR were exchanged into LS buffer (50 mM HEPES, pH 7.4) with specified NaCl concentration and centrifuged for 30 min at 100,000 rpm (TLA100, Beckman Coulter). Unless otherwise stated, 10 μM GUN4 or 2.7 μM GluTR was mixed with an equal volume of cpSRP43 at indicated concentrations in the same buffer at 4 °C. The chaperone-client mixtures were incubated in a water bath at 42 °C for 5 min and placed on ice for 5 min. The optical density at 360 nm was recorded, normalized to that of the client without cpSRP43, and plotted as a function of cpSRP43 concentration. The data were fit to Eq. 4 where possible. In cases where an increase in turbidity was observed at sub-stoichiometric cpSRP43 or where no protection was detectable, a smoothing spline was plotted and *K*_*1/2 sol*_, the cpSRP43 concentration at which 50% of client protein was solubilized, was extrapolated and used in lieu of *K*_*sol*_.

### GUN4 aggregation kinetics

Purified GUN4 was centrifuged for 30 min at 100,000 rpm (TLA100, Beckman Coulter) and diluted to 1-10 μM in LS buffer with indicated NaCl concentrations. Proteins were incubated in a water bath set between 30-46 °C for 1-60 minutes and then placed on ice for 5 minutes. The endpoint optical density at 360 nm (A_360nm_) was recorded. Kinetic data was fit to a single exponential function (Eq. 5),

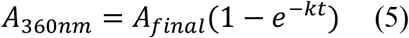

in which *A*_*final*_ is the optical density at 360 nm when the reaction reached equilibrium, *k* is the misfolding rate constant, and t is the incubation time at elevated temperature.

### Fluorescence anisotropy

Binding of cpSRP43 to HiLyte-Fluor488-labeled L11 peptide (GSFDPLGLADD) or to fluorescein-labeled 54C peptide (QKQKAPPGTARRKRKAC) was measured based on fluorescence anisotropy, as previously described (*10*). Measurements were performed in CD buffer with 50 mM NaCl at specified temperatures on a FluoroLog 3-22 spectrofluorometer (HORIBA), using 100 nM labeled peptide and the indicated concentrations of cpSRP43. The samples were excited at 500 nm, and fluorescence anisotropy was recorded at 527 nm. The observed anisotropy value (A_obs_) as a function of cpSRP43 concentration ([cpSRP43]) was fit to Eq. 6a,

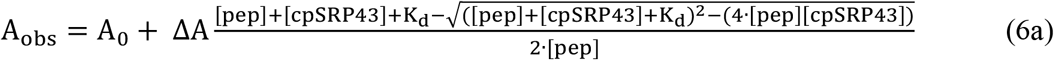

in which A_0_ is the anisotropy value of the peptide alone, ΔA is the change in anisotropy at saturating concentrations of cpSRP43, [pep] is peptide concentration, and *K*_*d*_ is the equilibrium dissociation constant for the interaction between cpSRP43 and peptide. Where noted, data are reported as the fraction bound, calculated for each A_obs_ based on Eq. 6b,

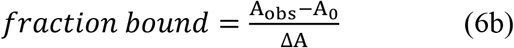

### BTFA labeling

cpSRP43 variants containing a single cysteine at residue 150 (D150C, C175A, C297S) were reduced with 4 mM DTT and exchanged into degassed labeling buffer (50 mM HEPES, 300 mM NaCl, 1 mM EDTA 10% glycerol, pH 7.4). 50 µM cpSRP43 was labeled with 2 mM BTFA (Sigma Aldrich) at room temperature for 2 hours. Labeled cpSRP43 was purified on a HiPrep (Cytiva) column in CD buffer with 50 mM NaCl, and 10% D_2_O was added to the final sample. The final concentrations of BTFA-labeled cpSRP43 were 35-60 µM.

### NMR spectroscopy

^19^F NMR spectra were acquired in CD buffer with 50 mM NaCl at the indicated temperatures on a Bruker Avance 600 spectrometer equipped with a 5 mm QCI 1H/19F/13C/15N quadruple resonance cryoprobe with a single axis Z-gradient. Where specified, urea or 54C or L18 (VDPLYPGGSFDPLGLADD) peptides were added at the indicated concentrations. NMR data were processed with NMRPipe (*44*). Peaks were analyzed in MestReNova with peak deconvolutions fit to a generalized Lorentzian using simulated annealing (*45*). Peak areas were used to determine the relative population of *closed* and *open* states.

TROSY HSQC spectra of ^2^H,^15^N-labeled cpSRP43 (WT, R189L or DE3NQ) were recorded on an 800 MHz Bruker Avance spectrometer equipped with a 5mm TCI 1H/13C/15N triple resonance cryoprobe with a single axis Z-gradient. NMR spectra were acquired at 17 °C in NMR buffer (50 mM Na_2_HPO_4_, pH 6.5, 150 mM NaCl) supplemented with 10% vol/vol D_2_O. NMR data were processed with NMRPipe (*44*). Data was visualized and analyzed with NMRviewJ (*46*). Assignments were transferred from Liang et al., 2016 (*10*) and spectra were normalized to the average intensity of the flexible N-terminus (1-30).

### Statistical Analysis

Gaussian (normal) distribution was used as the statistical model for error estimates. Statistics are described for all reported values in the figure legends.

## Acknowledgments

We thank members of the Shan group for helpful comments on the manuscript.

## Author contributions

Conceptualization: ARS, SS

Methodology: GK

Investigation: ARS, GK

Visualization: ARS

Supervision: SS, PW

Writing—original draft: ARS

Writing—review & editing: SS, ARS, PW

## Data and materials availability

All data supporting the conclusions are included in the article main text and Supplementary Information.

## References

1. P. Jarvis, E. López-Juez, Biogenesis and homeostasis of chloroplasts and other plastids. Nature Reviews Molecular Cell Biology 14, 787–802 (2013).

2. L. S. Leutwiler, E. M. Meyerowitz, E. M. Tobin, Structure and expression of three light-harvesting chlorophyll a/b-binding protein genes in Arabidopsis thaliana. Nucleic Acids Research 14, 4051–4064 (1986).

3. G. F. Plumley, G. W. Schmidt, Light-Harvesting Chlorophyll a/b Complexes: Interdependent Pigment Synthesis and Protein Assembly. The Plant Cell 7, 689–704 (1995).

4. L. Dall’Osto, M. Bressan, R. Bassi, Biogenesis of light harvesting proteins. Biochimica et Biophysica Acta (BBA) - Bioenergetics 1847, 861–871 (2015).

5. H. Paulsen, B. Finkenzeller, N. Kühlein, Pigments induce folding of light-harvesting chlorophyll a/b-binding protein. European Journal of Biochemistry 215, 809–816 (1993).

6. K. Apel, H. Hirt, Reactive oxygen species: metabolism, oxidative stress, and signal transduction. Annu Rev Plant Biol 55, 373–399 (2004).

7. P. Wang, F.-C. Liang, D. Wittmann, A. Siegel, S. Shan, B. Grimm, Chloroplast SRP43 acts as a chaperone for glutamyl-tRNA reductase, the rate-limiting enzyme in tetrapyrrole biosynthesis. Proceedings of the National Academy of Sciences 115, E3588–E3596 (2018).

8. P. Jaru-Ampornpan, K. Shen, V. Q. Lam, M. Ali, S. Doniach, T. Z. Jia, S. Shan, ATP-independent reversal of a membrane protein aggregate by a chloroplast SRP subunit. Nat Struct Mol Biol 17, 696–702 (2010).

9. K. F. Stengel, I. Holdermann, P. Cain, C. Robinson, K. Wild, I. Sinning, Structural Basis for Specific Substrate Recognition by the Chloroplast Signal Recognition Particle Protein cpSRP43. Science 321, 253–256 (2008).

10. F.-C. Liang, G. Kroon, C. Z. McAvoy, C. Chi, P. E. Wright, S. Shan, Conformational dynamics of a membrane protein chaperone enables spatially regulated substrate capture and release. Proceedings of the National Academy of Sciences 113, E1615–E1624 (2016).

11. P. Cain, I. Holdermann, I. Sinning, A. E. Johnson, C. Robinson, Binding of chloroplast signal recognition particle to a thylakoid membrane protein substrate in aqueous solution and delineation of the cpSRP43-substrate interaction domain. Biochem. J. 437, 149–155 (2011).

12. C. Z. McAvoy, A. Siegel, S. Piszkiewicz, E. Miaou, M. Yu, T. Nguyen, A. Moradian, M. J. Sweredoski, S. Hess, S. Shan, Two distinct sites of client protein interaction with the chaperone cpSRP43. Journal of Biological Chemistry 293, 8861–8873 (2018).

13. I. Holdermann, N. H. Meyer, A. Round, K. Wild, M. Sattler, I. Sinning, Chromodomains read the arginine code of post-translational targeting. Nature Structural & Molecular Biology 19, 260–263 (2012).

14. F. Gao, A. D. Kight, R. Henderson, S. Jayanthi, P. Patel, M. Murchison, P. Sharma, R. L. Goforth, T. K. S. Kumar, R. L. Henry, C. D. Heyes, Regulation of Structural Dynamics within a Signal Recognition Particle Promotes Binding of Protein Targeting Substrates. J. Biol. Chem. 290, 15462–15474 (2015).

15. K. M. Kathir, D. Rajalingam, V. Sivaraja, A. Kight, R. L. Goforth, C. Yu, R. Henry, T. K. S. Kumar, Assembly of Chloroplast Signal Recognition Particle Involves Structural Rearrangement in cpSRP43. Journal of Molecular Biology 381, 49–60 (2008).

16. C.-J. Tu, D. Schuenemann, N. E. Hoffman, Chloroplast FtsY, Chloroplast Signal Recognition Particle, and GTP Are Required to Reconstitute the Soluble Phase of Light-harvesting Chlorophyll Protein Transport into Thylakoid Membranes. J. Biol. Chem. 274, 27219–27224 (1999).

17. M. Moore, M. S. Harrison, E. C. Peterson, R. Henry, Chloroplast Oxa1p homolog albino3 is required for post-translational integration of the light harvesting chlorophyll-binding protein into thylakoid membranes. J. Biol. Chem. 275, 1529–1532 (2000).

18. S. Falk, S. Ravaud, J. Koch, I. Sinning, The C terminus of the Alb3 membrane insertase recruits cpSRP43 to the thylakoid membrane. J. Biol. Chem. 285, 5954–5962 (2010).

19. B. Dünschede, T. Bals, S. Funke, D. Schünemann, Interaction studies between the chloroplast signal recognition particle subunit cpSRP43 and the full-length translocase Alb3 reveal a membrane-embedded binding region in Alb3 protein. J. Biol. Chem. 286, 35187–35195 (2011).

20. N. E. Lewis, N. J. Marty, K. M. Kathir, D. Rajalingam, A. D. Kight, A. Daily, T. K. S. Kumar, R. L. Henry, R. L. Goforth, A Dynamic cpSRP43-Albino3 Interaction Mediates Translocase Regulation of Chloroplast Signal Recognition Particle (cpSRP)-targeting Components. J. Biol. Chem. 285, 34220–34230 (2010).

21. L. A. Eichacker, R. Henry, Function of a chloroplast SRP in thylakoid protein export. Biochimica et Biophysica Acta (BBA) - Molecular Cell Research 1541, 120–134 (2001).

22. D. Schuenemann, S. Gupta, F. Persello-Cartieaux, V. I. Klimyuk, J. D. G. Jones, L. Nussaume, N. E. Hoffman, A novel signal recognition particle targets light-harvesting proteins to the thylakoid membranes. Proceedings of the National Academy of Sciences 95, 10312–10316 (1998).

23. V. I. Klimyuk, F. Persello-Cartieaux, M. Havaux, P. Contard-David, D. Schuenemann, K. Meiherhoff, P. Gouet, J. D. Jones, N. E. Hoffman, L. Nussaume, A chromodomain protein encoded by the arabidopsis CAO gene is a plant-specific component of the chloroplast signal recognition particle pathway that is involved in LHCP targeting. Plant Cell 11, 87–99 (1999).

24. X. Li, R. Henry, J. Yuan, K. Cline, N. E. Hoffman, A chloroplast homologue of the signal recognition particle subunit SRP54 is involved in the posttranslational integration of a protein into thylakoid membranes. PNAS 92, 3789–3793 (1995).

25. T. Tzvetkova-Chevolleau, C. Hutin, L. D. Noël, R. Goforth, J.-P. Carde, S. Caffarri, I. Sinning, M. Groves, J.-M. Teulon, N. E. Hoffman, R. Henry, M. Havaux, L. Nussaume, Canonical Signal Recognition Particle Components Can Be Bypassed for Posttranslational Protein Targeting in Chloroplasts. The Plant Cell 19, 1635–1648 (2007).

26. S. Ji, A. Siegel, S. Shan, B. Grimm, P. Wang, Chloroplast SRP43 autonomously protects chlorophyll biosynthesis proteins against heat shock. Nat. Plants, 1–13 (2021).

27. A. Siegel, C. Z. McAvoy, V. Lam, F.-C. Liang, G. Kroon, E. Miaou, P. Griffin, P. E. Wright, S.-O. Shan, A Disorder-to-Order Transition Activates an ATP-Independent Membrane Protein Chaperone. J Mol Biol 432, 166708 (2020).

28. N. J. Greenfield, Using circular dichroism spectra to estimate protein secondary structure. Nat Protoc 1, 2876–90 (2006).

29. K. K. Lee, C. A. Fitch, B. García-Moreno E., Distance dependence and salt sensitivity of pairwise, coulombic interactions in a protein. Protein Science 11, 1004–1016 (2002).

30. P. Jaru-Ampornpan, F.-C. Liang, A. Nisthal, T. X. Nguyen, P. Wang, K. Shen, S. L. Mayo, S. Shan, Mechanism of an ATP-independent Protein Disaggregase: II. Distinct Molecular Interactions Drive Multiple Steps During Aggregate Assembly. Journal of Biological Chemistry 288, 13431–13445 (2013).

31. C. J. Tu, E. C. Peterson, R. Henry, N. E. Hoffman, The L18 Domain of Light-harvesting Chlorophyll Proteins Binds to Chloroplast Signal Recognition Particle 43. J. Biol. Chem. 275, 13187–13190 (2000).

32. T. M. Franzmann, P. Menhorn, S. Walter, J. Buchner, Activation of the Chaperone Hsp26 Is Controlled by the Rearrangement of Its Thermosensor Domain. Molecular Cell 29, 207–216 (2008).

33. F. Stengel, A. J. Baldwin, A. J. Painter, N. Jaya, E. Basha, L. E. Kay, E. Vierling, C. V. Robinson, J. L. P. Benesch, Quaternary dynamics and plasticity underlie small heat shock protein chaperone function. Proceedings of the National Academy of Sciences 107, 2007–2012 (2010).

34. L. Foit, J. S. George, B. W. Zhang, C. L. Brooks, J. C. A. Bardwell, Chaperone activation by unfolding. Proceedings of the National Academy of Sciences 110, E1254–E1262 (2013).

35. P. Rajagopal, E. Tse, A. J. Borst, S. P. Delbecq, L. Shi, D. R. Southworth, R. E. Klevit, A conserved histidine modulates HSPB5 structure to trigger chaperone activity in response to stress-related acidosis. eLife 4, e07304 (2015).

36. A. F. Clouser, R. E. Klevit, pH-dependent structural modulation is conserved in the human small heat shock protein HSBP1. Cell Stress and Chaperones 22, 569–575 (2017).

37. T. Fleckenstein, A. Kastenmüller, M. L. Stein, C. Peters, M. Daake, M. Krause, D. Weinfurtner, M. Haslbeck, S. Weinkauf, M. Groll, J. Buchner, The Chaperone Activity of the Developmental Small Heat Shock Protein Sip1 Is Regulated by pH-Dependent Conformational Changes. Molecular Cell 58, 1067–1078 (2015).

38. B. Groitl, S. Horowitz, K. A. T. Makepeace, E. V. Petrotchenko, C. H. Borchers, D. Reichmann, J. C. A. Bardwell, U. Jakob, Protein unfolding as a switch from self-recognition to high-affinity client binding. Nat Commun 7, 10357 (2016).

39. J. Winter, M. Ilbert, P. C. F. Graf, D. Özcelik, U. Jakob, Bleach Activates a Redox-Regulated Chaperone by Oxidative Protein Unfolding. Cell 135, 691–701 (2008).

40. T. Krojer, J. Sawa, E. Schäfer, H. R. Saibil, M. Ehrmann, T. Clausen, Structural basis for the regulated protease and chaperone function of DegP. Nature 453, 885–890 (2008).

41. D. Zurawa-Janicka, M. Jarzab, A. Polit, J. Skorko-Glonek, A. Lesner, A. Gitlin, A. Gieldon, J. Ciarkowski, P. Glaza, A. Lubomska, B. Lipinska, Temperature-induced changes of HtrA2(Omi) protease activity and structure. Cell Stress and Chaperones 18, 35–51 (2013).

42. Y. Toyama, R. W. Harkness, L. E. Kay, Structural basis of protein substrate processing by human mitochondrial high-temperature requirement A2 protease. Proceedings of the National Academy of Sciences 119, e2203172119 (2022).

43. P. Jaru-Ampornpan, S. Chandrasekar, S. Shan, Efficient Interaction between Two GTPases Allows the Chloroplast SRP Pathway to Bypass the Requirement for an SRP RNA. Molecular Biology of the Cell 18, 2636–2645 (2007).

44. F. Delaglio, S. Grzesiek, G. W. Vuister, G. Zhu, J. Pfeifer, A. Bax, NMRPipe: A multidimensional spectral processing system based on UNIX pipes. Journal of Biomolecular NMR 6, 277–293 (1995).

45. M. R. Willcott, MestRe Nova. J. Am. Chem. Soc. 131, 13180–13180 (2009).

46. B. A. Johnson, R. A. Blevins, NMR View: A computer program for the visualization and analysis of NMR data. J. Biomol. NMR 4, 603–614 (1994).

