## Supplementary Information for "Switchable client specificity in a dual functional chaperone coordinates light harvesting complex biogenesis"

**This PDF file includes:**

Figs. S1 to S9
Table S1

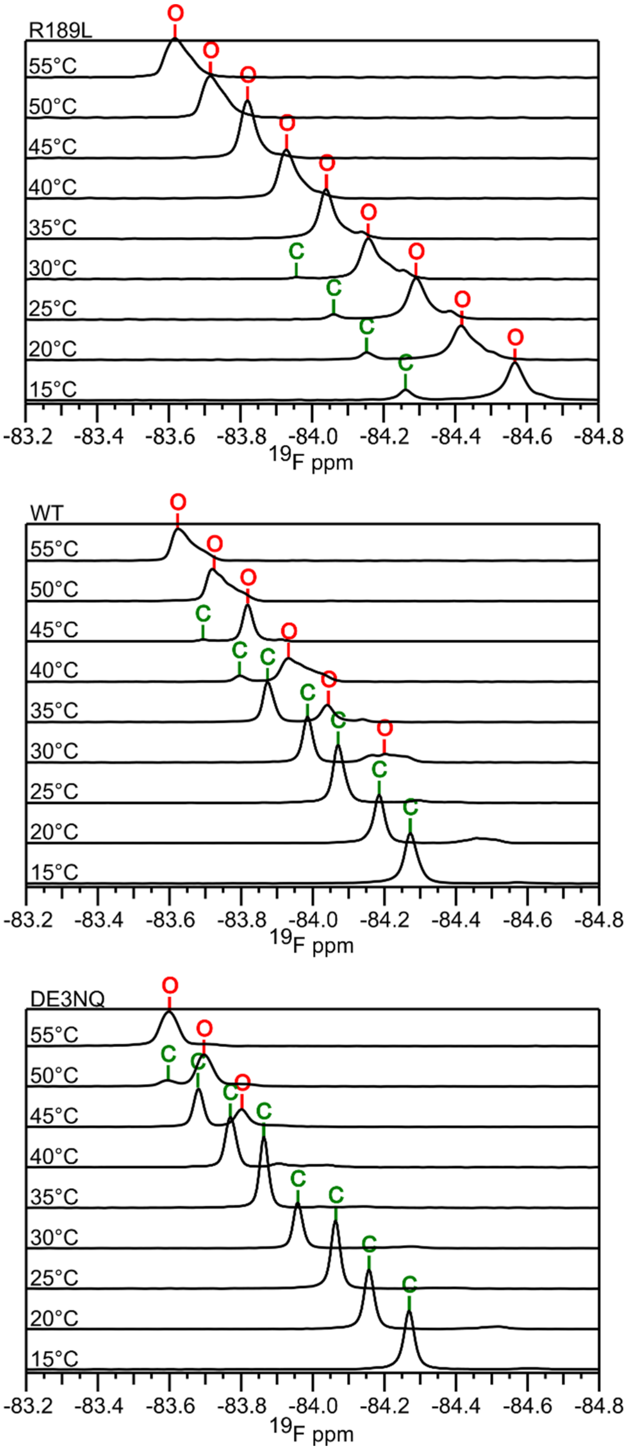


**Figure S1. $^{19}$F-NMR spectra for BTFA-labeled cpSRP43 variants at rising temperature.** Data were collected as in Fig. 1A. Note that the chemical shift of 19F-BTFA moves downfield at higher temperatures, but this does not interfere with data interpretation. WT spectra from Fig. 1A are reproduced for comparison.

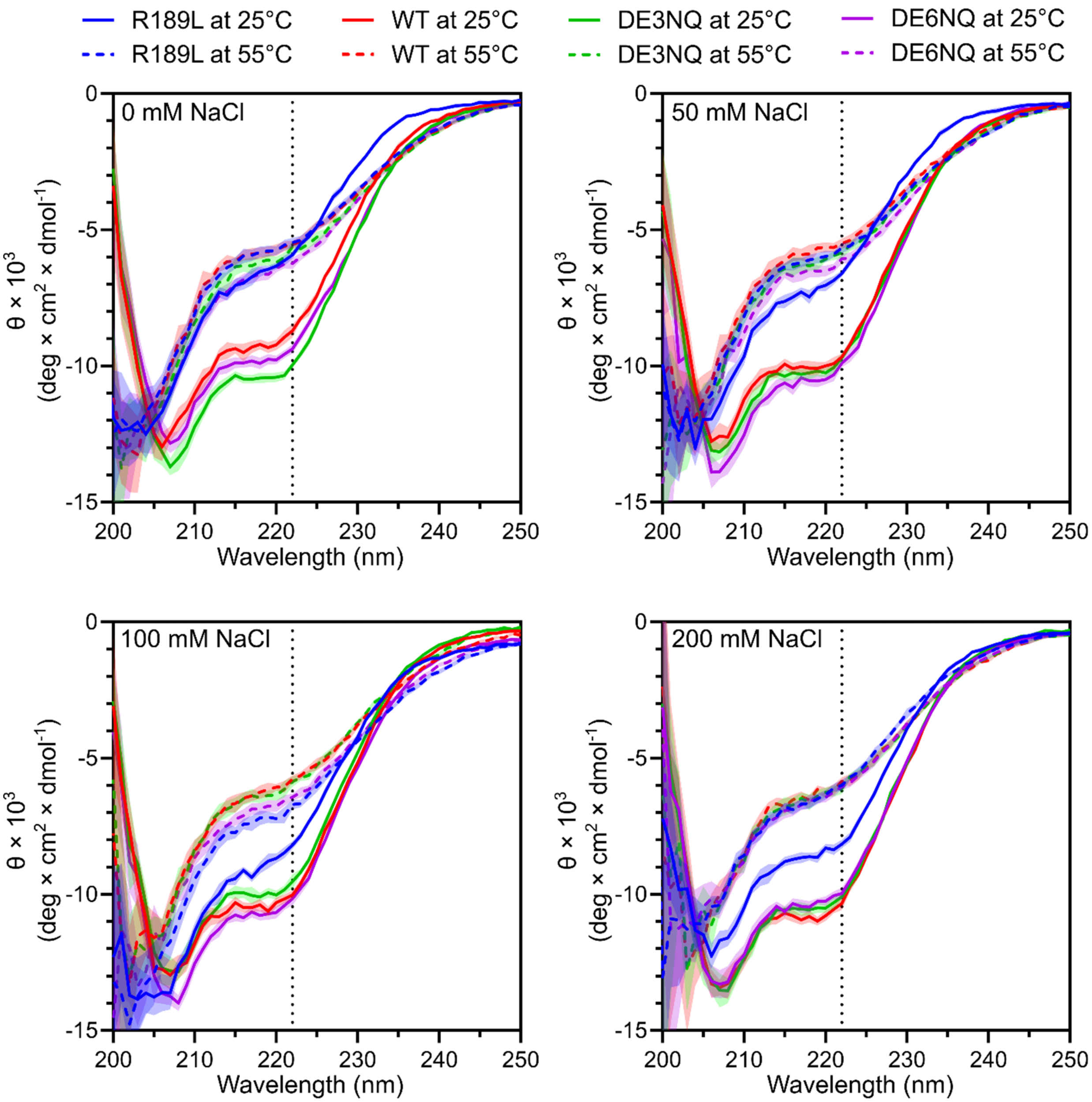


**Figure S2. CD spectra of cpSRP43 variants at different ionic strength.** Data were collected with 10 μM of each cpSRP43 variant in CD buffer at the indicated salt concentrations at 25 °C (solid lines) and 55 °C (dotted lines). Values represent mean residue ellipticity ± S.D. over 5 seconds. The dotted line at 222 nm marks the peak that reports primarily on α-helical content and was used throughout CD thermal melts. Values below 205 nm were affected by $Cl^-$ ions leading to high error and low confidence.

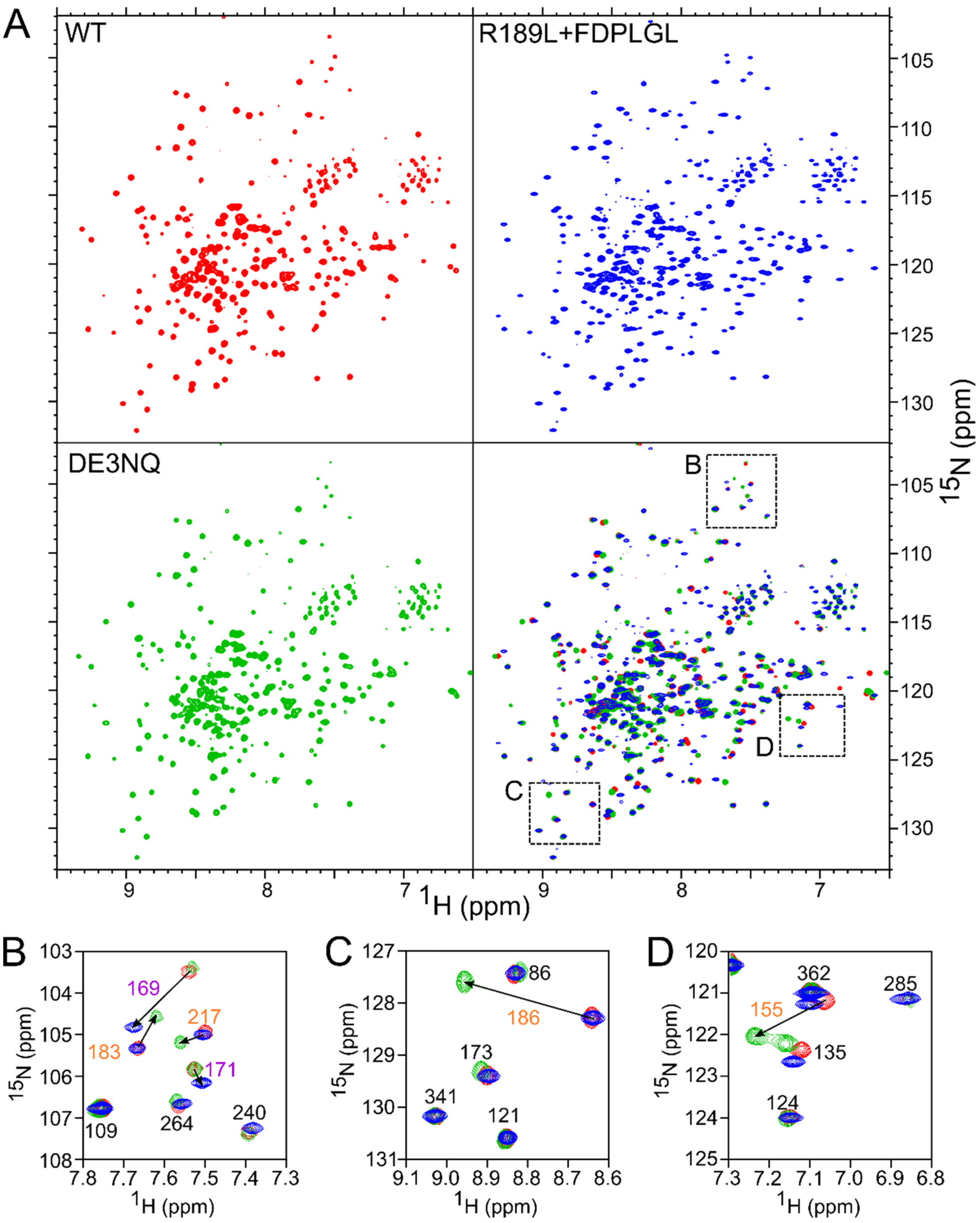


**Figure S3. HSQC NMR spectra of cpSRP43 variants**. (**A**) $^{1}$H, $^{15}$N-TROSY spectra of WT cpSRP43 (red), DE3NQ (green), R189L bound with the L18 peptide (blue), and all three overlaid (bottom right). (**B-D**) Zoom in of three regions from (A). Assigned residues (from Liang et al, 2016.[1]) labeled in *orange* are within 10 Å of DE3NQ, and those in *violet* are within 10 Å of the FDPLGL motif binding site (based on the *closed* state structure). Arrows indicate peak shifts relative to WT.

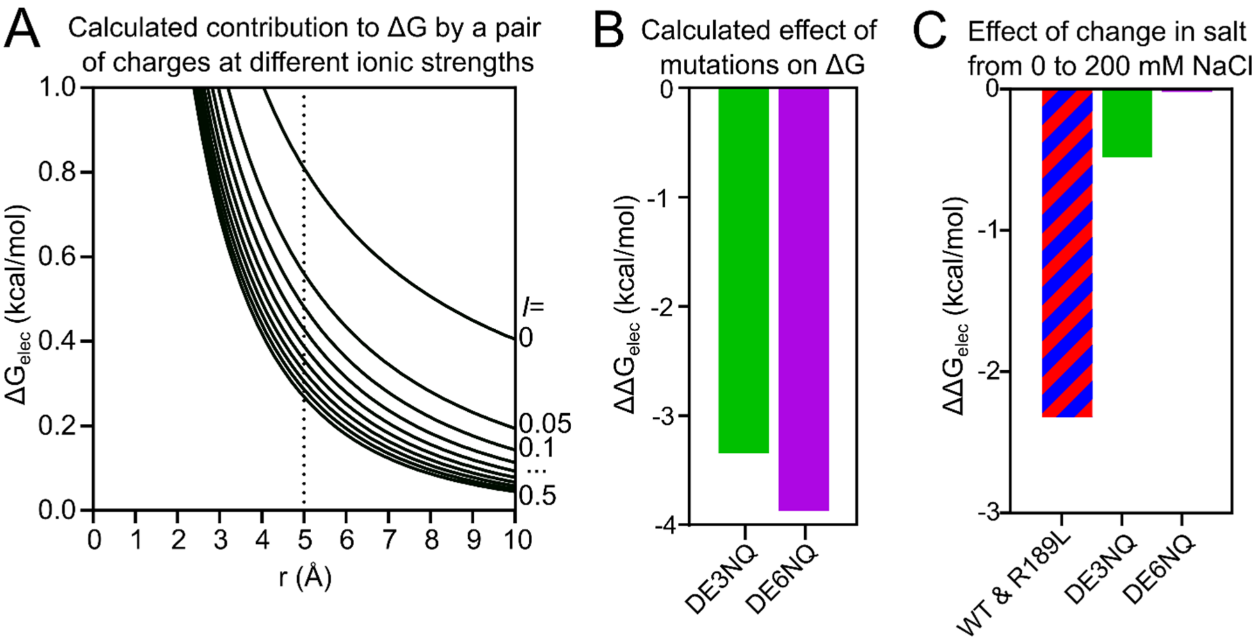


**Figure S4. Calculated effect of ionic strength and mutations on the stability of the closed state of cpSRP43.** (**A**) Free energy contributions from Coulomb effects ($\Delta G_{electric}$) of a charged pair separated by radius (r), calculated using Eq. 3 at varying ionic strengths (I). Dotted line at 5 Å marks the distance between neighboring acidic residues in cpSRP43. (**B**) Predicted changes in $\Delta G_{electric}$ ($\Delta\Delta G_{elec}$) caused by the DE3NQ and DE6NQ mutations in 50 mM NaCl buffer (I = 0.1) based on the distances between neighboring charges across the 7 acidic residues in the *closed* state structure. (**C**) Calculation of the contribution of the 7 acidic residues to $\Delta G_{elec}$ when the NaCl concentration is increased from 0 mM (I = 0.05) to 200 mM (I = 0.25) in each cpSRP43 variant.

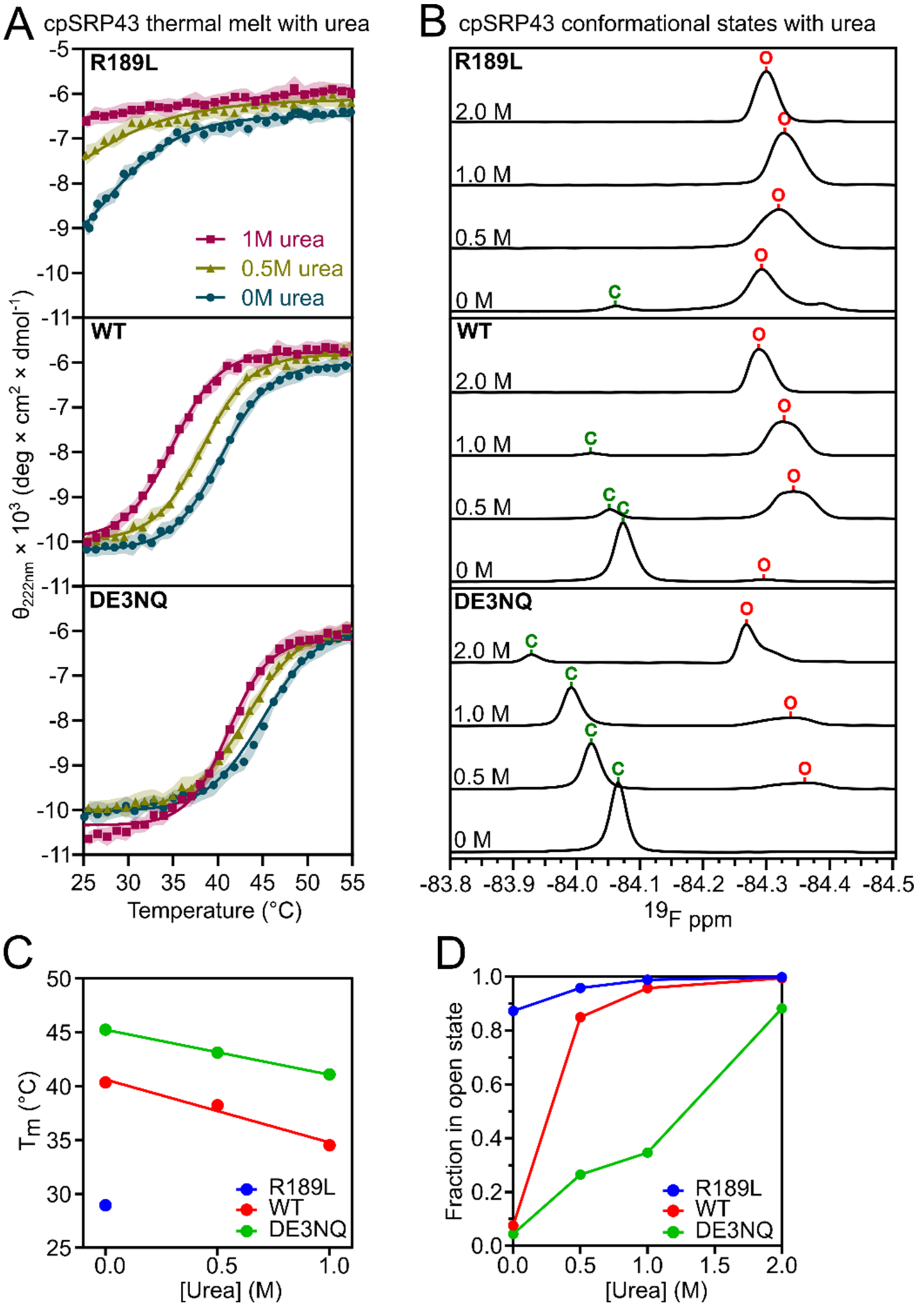


**Figure S5. Low doses of urea drive the opening of R189L and WT cpSRP43, but not DE3NQ.** (**A**) Effect of urea on the helical content of R189L, WT, and DE3NQ cpSRP43, measured by molar ellipticity at 222 nm in CD Buffer with 200 mM NaCl and 0/0.5/1 M urea. Values represent mean residue ellipticity ± S.D. over 10 seconds The lines are fits of the data to Eq. 2, and the obtained $T_m$ values are summarized in Table 1. (**B**) Effect of urea on the $^{19}$F NMR spectra of cpSRP43 variants, measured in CD Buffer with 50 mM NaCl at 25 °C. The *closed* and *open* state peaks are marked. (**C-D**) Effect of urea on the $T_m$ values (C; based on the measurements in A) and *open* state fraction (D; based on the data in B) of the cpSRP43 variants.

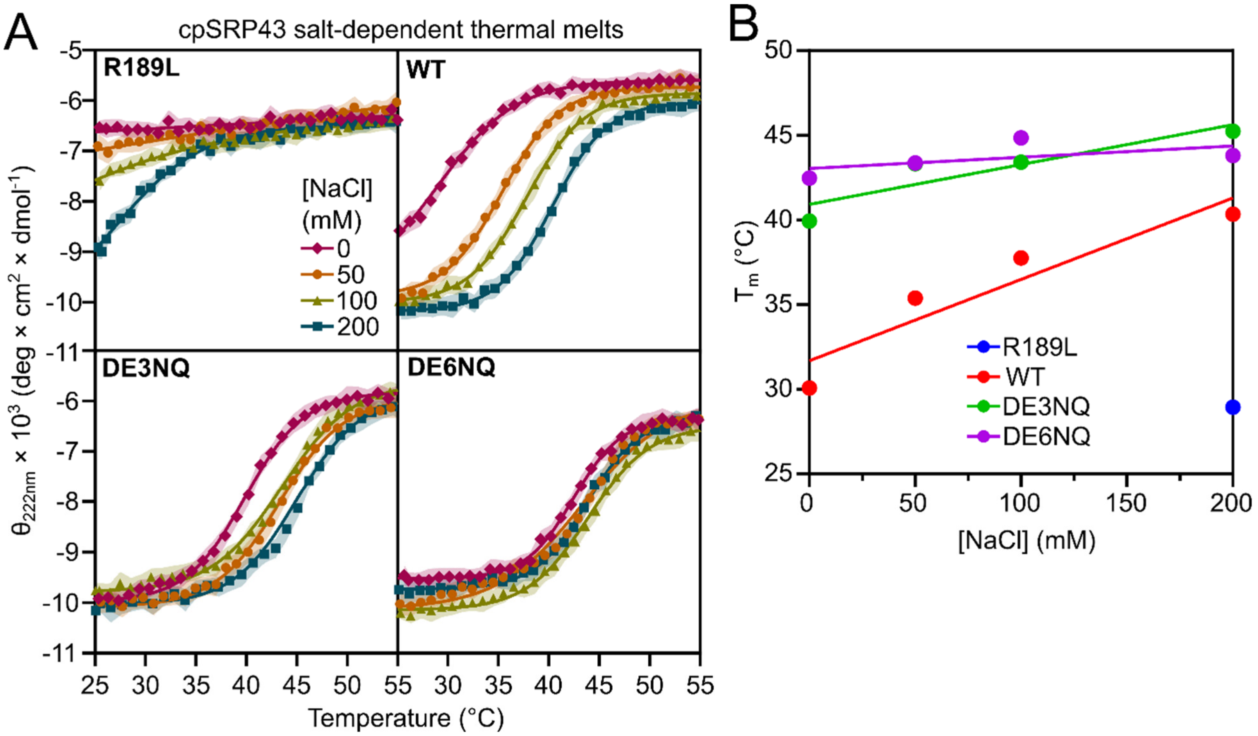


**Figure S6. Higher ionic strength stabilizes closed cpSRP43.** (**A**) Heat-induced opening of cpSRP43 variants in CD Buffer with the indicated NaCl concentrations, measured using CD. Values represent mean residue ellipticity ± S.D. over 10 seconds. The lines are fits of the data to Eq. 2, and the obtained $T_m$ values are summarized in Table 1. (**B**) The $T_m$ values obtained from (A) as a function of salt concentration.

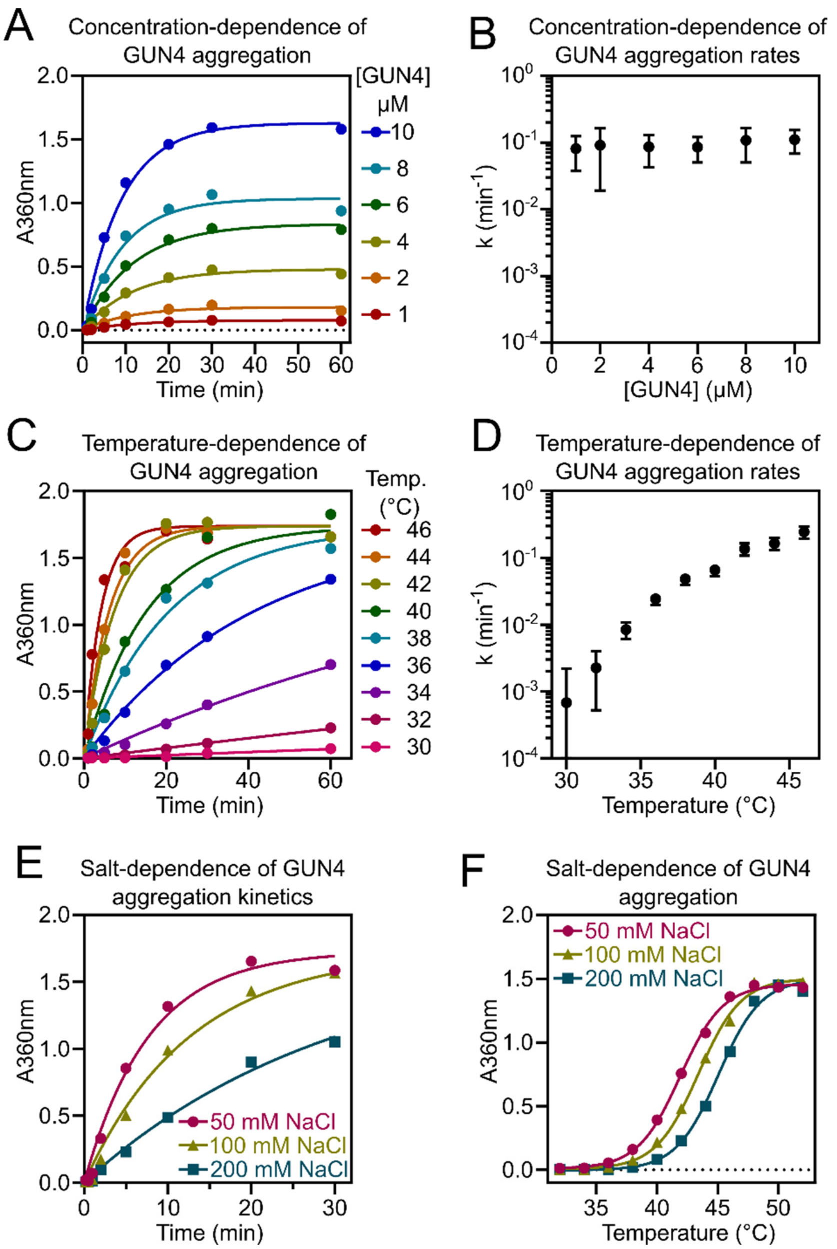


**Figure S7. GUN4 aggregation is rate-limited by misfolding and rises with higher temperature and lower salt.** (**A, C, E**) Time courses of heat-induced aggregation of GUN4 at the indicated GUN4 concentrations (A), temperatures (C), and salt concentrations (E). Reactions were carried out with 10 µM GUN4 at 42 °C and 50 mM NaCl unless otherwise specified. The data were fit to Eq. 5, and the obtained aggregation rate constants (*k*) and optical density at equilibrium ($A_{360,\ final}$) are summarized in Table S1. (**B, D**) GUN4 aggregation kinetics is unchanged with different GUN4 concentration (B), but rises with increasing temperature (D). Values represent fitted value ± 95% CI of the fit. (**F**) Heat-induced aggregation of GUN4 at different salt concentrations, based on $A_{360}$ reading at 5 minutes.

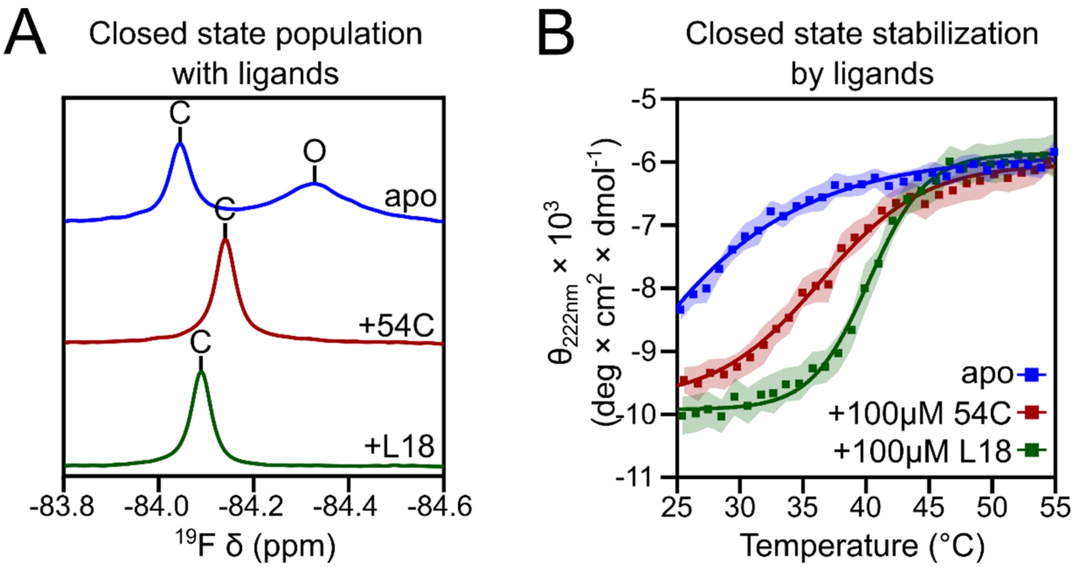


**Figure S8. Ligand binding stabilizes the closed state of cpSRP43.** (**A**) $^{19}$F NMR spectra of BTFA-labeled R189L in the apo (blue), 54C-bound (red), and L18-bound (green) states. Data were collected at 17 °C in NMR Buffer. (**B**) CD thermal melts monitoring the *closed*-to-*open* transition of apo (blue), 54C-bound (red), and L18-bound (green) R189L. Data were collected in CD buffer with 200 mM NaCl. Values represent mean residue ellipticity ± S.D. over 10 seconds. The lines are fits of the data to Eq. 2.

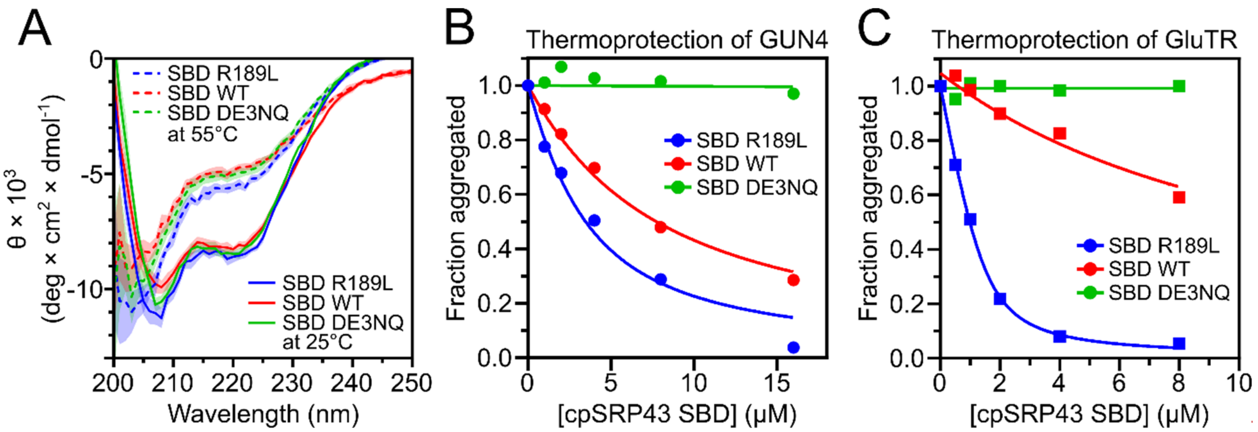


**Figure S9. cpSRP43 SBD stability and thermoprotection of GUN4 and GluTR.** (**A**) CD spectra of cpSRP43(SBD) with the indicated mutations. Values represent mean residue ellipticity ± S.D. over 5 seconds. (**B, C**) Turbidity assays to measure the protection of GUN4 (B) or GluTR (C) by the SBD of the indicated chaperone variants. Measurements were carried out at 42 °C for 5 minutes.

**Table S1. Summary of the observed rate of GUN4 aggregation (*k*) and the optical density at 360 nm ($A_{360,\ final}$) when the reaction reached equilibrium.**

**A.** Fitted parameters ± S.E. from the fit for data in Figure S7A.

| **[GUN4] (µM)** | **1** | **2** | **4** | **6** | **8** | **10** |
|---|---|---|---|---|---|---|
| $k$ ($min^{-1}$) | 0.081 ±0.02 | 0.091 ±0.03 | 0.086 ±0.02 | 0.085 ±0.01 | 0.11 ±0.02 | 0.11 ±0.02 |
| $A_{360,\ final}$ | 0.078 ±0.006 | 0.18 ±0.02 | 0.478 ±0.03 | 0.83 ±0.05 | 1.03 ±0.07 | 1.63 ±0.08 |

**B.** Fitted parameters ± S.E. from the fit for data in Figure S7C.

| **T (°C)** | **30** | **32** | **34** | **36** | **38** | **40** | **42** | **44** | **46** |
|---|---|---|---|---|---|---|---|---|---|
| $k$ ($min^{-1}$) | 0.00068 ±0.0008 | 0.0023 ±0.0008 | 0.0084 ±0.001 | 0.024 ±0.002 | 0.048 ±0.004 | 0.066 ±0.006 | 0.13 ±0.01 | 0.16 ±0.02 | 0.24 ±0.02 |
| $A_{360,\ final}$* | 1.74 ± 0.03 | 1.74 ± 0.03 | 1.74 ± 0.03 | 1.74 ± 0.03 | 1.74 ± 0.03 | 1.74 ± 0.03 | 1.74 ± 0.03 | 1.74 ± 0.03 | 1.74 ± 0.03 |

* The $A_{360,\ final}$ was fit as a shared value across all temperatures.

**C.** Fitted parameters ± S.E. from the fit for data in Figure S7E.

| **[NaCl] (mM)** | **50** | **100** | **200** |
|---|---|---|---|
| $k$ ($min^{-1}$) | 0.13 ± 0.01 | 0.079 ± 0.007 | 0.033 ± 0.003 |
| $A_{360,\ final}$* | 1.73 ± 0.06 | 1.73 ± 0.06 | 1.73 ± 0.06 |

* The $A_{360,\ final}$ was fit as a shared value across all salt conditions.
